# A genomic study of the Japanese population focusing on the glucocorticoid receptor interactome highlights distinct genetic characteristics associated with stress response

**DOI:** 10.1101/2022.09.16.508283

**Authors:** Thanasis Mitsis, Louis Papageorgiou, Eleni Papakonstantinou, Io Diakou, Katerina Pierouli, Konstantina Dragoumani, Flora Bacopoulou, Tomoshige Kino, George P Chrousos, Elias Eliopoulos, Dimitrios Vlachakis

## Abstract

All living organisms have been programmed to maintain a complex inner equilibrium called homeostasis, despite numerous adversities during their lifespan. Any threatening or perceived as such stimuli for homeostasis is termed a stressor, and a highly conserved response system called the stress response system has been developed to cope with these stimuli and maintain or reinstate homeostasis. The glucocorticoid receptor, a transcription factor belonging to the nuclear receptors protein superfamily, has a major role in the stress response system, and research on its’ interactome may provide novel information regarding the mechanisms underlying homeostasis maintenance. A list of 149 autosomal genes which have an essential role in GR function or are prime examples of GRE-containing genes was composed in order to gain a comprehensive view of the GR interactome. A search for SNPs on those particular genes was conducted on a dataset of 3.554 Japanese individuals, with mentioned polymorphisms being annotated with relevant information from the ClinVar, LitVar, and dbSNP databases. Forty-two SNPs of interest and their genomic locations were identified. These SNPs have been associated with drug metabolism and neuropsychiatric, metabolic, and immune system disorders, while most of them were located in intronic regions. The frequencies of those SNPs were later compared with a dataset consisting of 1465 Korean individuals in order to find population-specific characteristics based on some of the identified SNPs of interest. The results highlighted that rs1043618 frequencies were different in the two populations, with mentioned polymorphism having a potential role in chronic obstructive pulmonary disease in response to environmental stressors. This SNP is located in the HSPA1A gene which codes for an essential GR co-chaperone, and such information showcases that similar gene may be novel genomic targets for managing or combatting stress-related pathologies.

## Introduction

Living organisms maintain a dynamic inner equilibrium, both physiological and psychological, termed homeostasis. This equilibrium is continuously challenged by internal or external adverse forces termed stressors (1). The term stress refers to a state of threatened or perceived as such homeostasis (2). Since living organisms need to cope with numerous stressors during their lifespan, they have developed an intricate neuroendocrine system that includes both physiological and behavioral responses, called the stress response system. This system consists of the hypothalamic-pituitary-adrenal (HPA) axis and the locus coeruleus (LC)/noradrenaline (NE)-autonomic nervous system and features both central and peripheral components (Figure 1) (3). The central components of the stress system, located in the hypothalamus and brainstem, include: a) parvocellular neurons that release corticotropin-releasing hormone (CRH), b) paraventricular nuclei (PVN) neurons that release arginine vasopressin (AVP), c) CRH neurons of the paragigantocellular and parabranchial nuclei of the medulla and LC and d), norepinephrine (NE) cell groups in the pons and medulla, known comprising the LC/NE system. The stress system’s peripheral components include a) the peripheral part of the Hypothalamic-Pituitary-Adrenal (HPA) axis, b) components of the parasympathetic system and c) the efferent sympathetic adrenomedullary system (SAM) (2).

**Figure 1.**
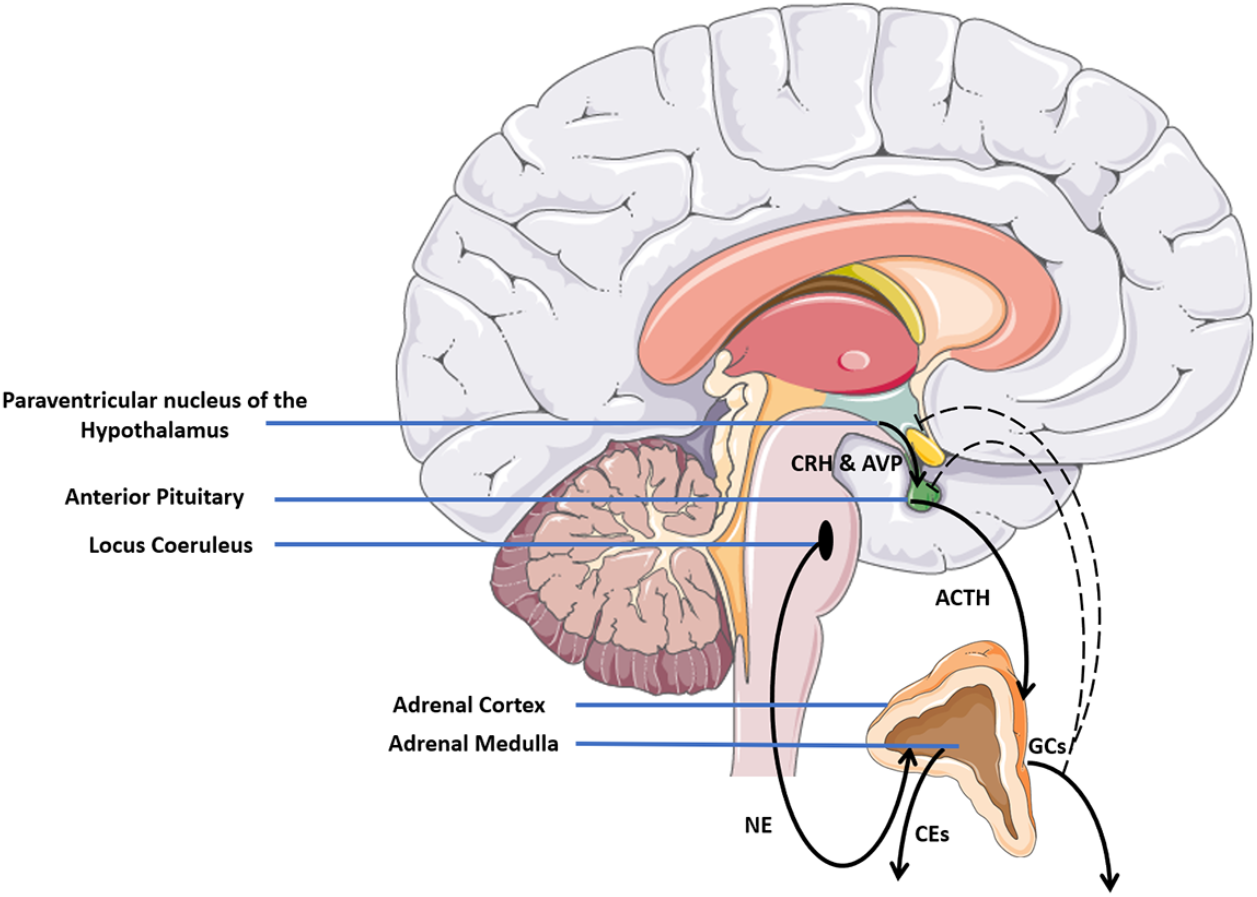
A schematic representation of the stress response system.

Specifically, in the HPA, CRH and AVP are secreted by the hypothalamus and act on the pituitary gland to trigger the adrenocorticotropic hormone production (ACTH). ACTH is then released into the circulation, which in turn stimulates glucocorticoid production and release from the adrenal cortex (4). Glucocorticoids, cortisol in humans and corticosterone in rodents are steroid hormones synthesized and released by the adrenal glands in a circadian manner (5–7). Cortisol can control CRH, AVP, and ACTH secretion in order to avoid prolonged HPA activity through sensitive negative feedback (8, 9). Under normal circumstances, the circadian rhythm followed by the HPA axis is characterized by high cortisol levels in the morning and low levels at night (10). The LC/NE and SAM systems, on the other hand, are mainly regulated by catecholamines (CEs). The locus coeruleus consists of a cluster of norepinephrine-producing neurons that are located in the upper dorsolateral pontine tegmentum and showcase branched axons, which project all through the neuraxis. These neurons are the sole source of NE to several brain regions, such as the hippocampus, neocortex, and cerebellum and can regulate the SAM system, which includes the NE neurons of the sympathetic system and the adrenal medulla (11, 12). Lastly, adrenal medulla stimulation by the LC/NE system leads to catecholamines secretion, specifically epinephrine (E) and norepinephrine (NE) which have a major role in the fight or flight response to stressors (3, 13).

Stress has the ability to alter gene expression through several mechanisms, including the direct effects of glucocorticoids (GCs), which are the final product of the HPA axis on gene transcription, plus activation of epigenetic mechanisms such as histone modifications and methylation/hydroxy-methylation of CpG residues in DNA (14). The biological processes influenced by such alterations include metabolism, development, reproduction, immune system pathways, and various cognitive functions (14). Thus, research on stress, the stress response system, homeostasis, glucocorticoids, and epigenetic modifications could provide both valuable information regarding human biological functions and possibly help develop medical applications.

Glucocorticoids act through binding with the high-affinity mineralocorticoid receptor (MR) and the low-affinity glucocorticoid receptor (GR), with GC action occurring mainly through the activation of the latter (9, 15). GR is almost exclusively activated by glucocorticoids, while MR can bind GCs and the mineralocorticoid aldosterone with similar high affinity (16). In the brain, a crucial component of the stress response system, MR is occupied at basal hormone levels due to its’ high affinity, while GR is activated at the circadian peak of glucocorticoid secretion and during stress (17). Thus, a research focus on GR can illuminate various aspects of the stress response system.

GR and MR are transcription factors that belong to the superfamily of nuclear receptors (6, 7, 9, 18). Nuclear receptors are ligand-dependent and regulate gene expression through DNA binding (19). The unliganded GR is predominantly localized within the cytoplasm, while ligand binding leads to the ligand-receptor complex’s translocation to the nucleus via the microtube network (20). In the absence of a ligand, GR is part of a protein complex with co-chaperones such as heat shock protein 70 (Hsp70), heat shock protein 40 (Hsp40), and heat shock protein 90 (Hsp90). After ligand binding, GR dissociates from its’ chaperone proteins and translocate to the nucleus in order to regulate gene transcription (21). More specifically, after receptor translation, GR forms a complex with Hsp40 and Hsp70. Following an ATP-dependent event, the Hsp40-Hsp70-GR complex is recruited by the Hsp70-Hsp90 Organizing Protein (Hop) to interact with Hsp90. After Hsp90 binding and another ATP-dependent event, Hop, Hsp40, and Hsp70 are dislodged from the chaperone complex and replaced by prostaglandin E synthase 3 (p23) and FK506 binding protein 51 (FKPB51). This specific GR complex showcases a high affinity for cortisol. Cortisol binding leads to conformational changes on GR and the replacement of FKPB51 by FK506 binding protein 52 (FKBP52), which lead to the receptor’s nuclear translocation (Figure 2) (22). GR induces transcription mainly by binding of GR homodimers to promoter regions that contain palindromic glucocorticoid response elements (GREs), a mechanism termed GR-dependent transactivation, while an alternate mechanism features the glucocorticoid receptor acting as a monomer and co-operating with other transcription factors (23, 24). After GR binding to GREs, the receptor acts as a scaffold for the assembly of several macromolecular complexes, which include coactivator proteins, chromatin remodeling factors, and mediators of the transcriptional machinery (20). GR-dependent transrepression, on the other hand, takes place mainly through GR interaction with DNA-bound transcription factors, while an alternate mechanism of transrepression is more similar to transactivation since GR binds to DNA sequences distinct from GREs, called negative GRE sites (nGREs) (20, 25).

**Figure 2.**
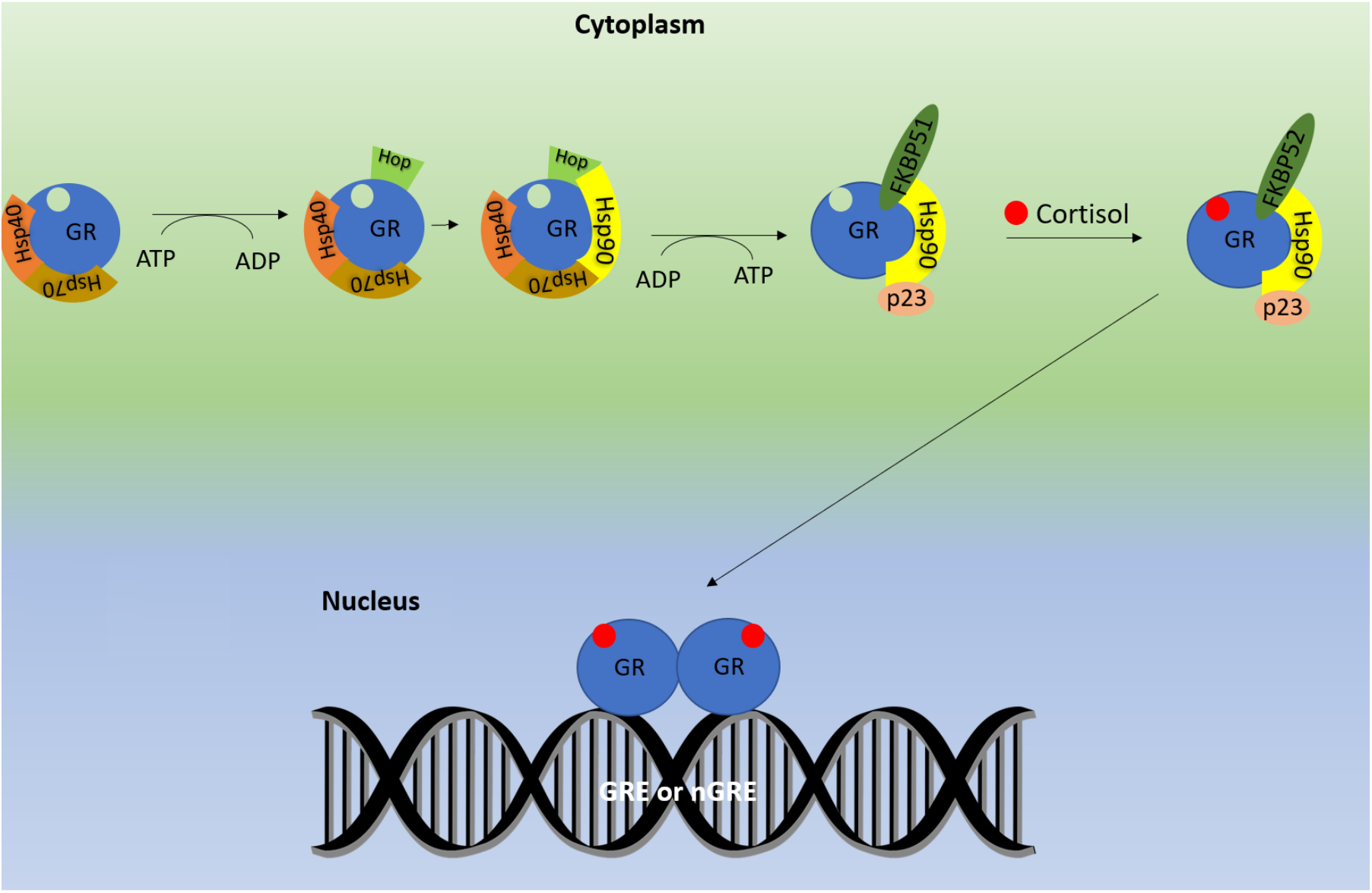
A schematic representation of GR signaling in gene regulation, specifically transactivation.

GR’s action is characteristic of its’ function in the stress response and highlights the importance of nuclear receptors interplay in biological functions. Regarding the latter, it is also important to note that GR has been shown to physically interact with MR and influence the action of other nuclear receptors such as estrogen receptor alpha, androgen receptor, retinoic acid receptors, and vitamin D receptor (26, 27). Thus, GR can also be used as a stepping-stone for providing insights into the complex interplay of nuclear receptor transcriptional networks and their contribution to homeostasis maintenance.

This study focuses on GR and genes that have an essential role in GR function or are prime examples of GR target genes. A distinct pipeline was followed to extract information in a precise and efficient way (Figure 3). A unique dataset consisting of single nuclear variations found in the autosomes of 3.554 Japanese individuals was used. The dataset was analyzed towards to finding (Single Nucleotide Polymorphisms) SNPs in the aforementioned genes, and if mentioned polymorphisms have been associated with human physiopathology. The results were then compared to a dataset featuring Koran individuals to find characteristics unique to the current population and if mentioned, characteristics can be associated with homeostasis mechanisms.

**Figure 3.**
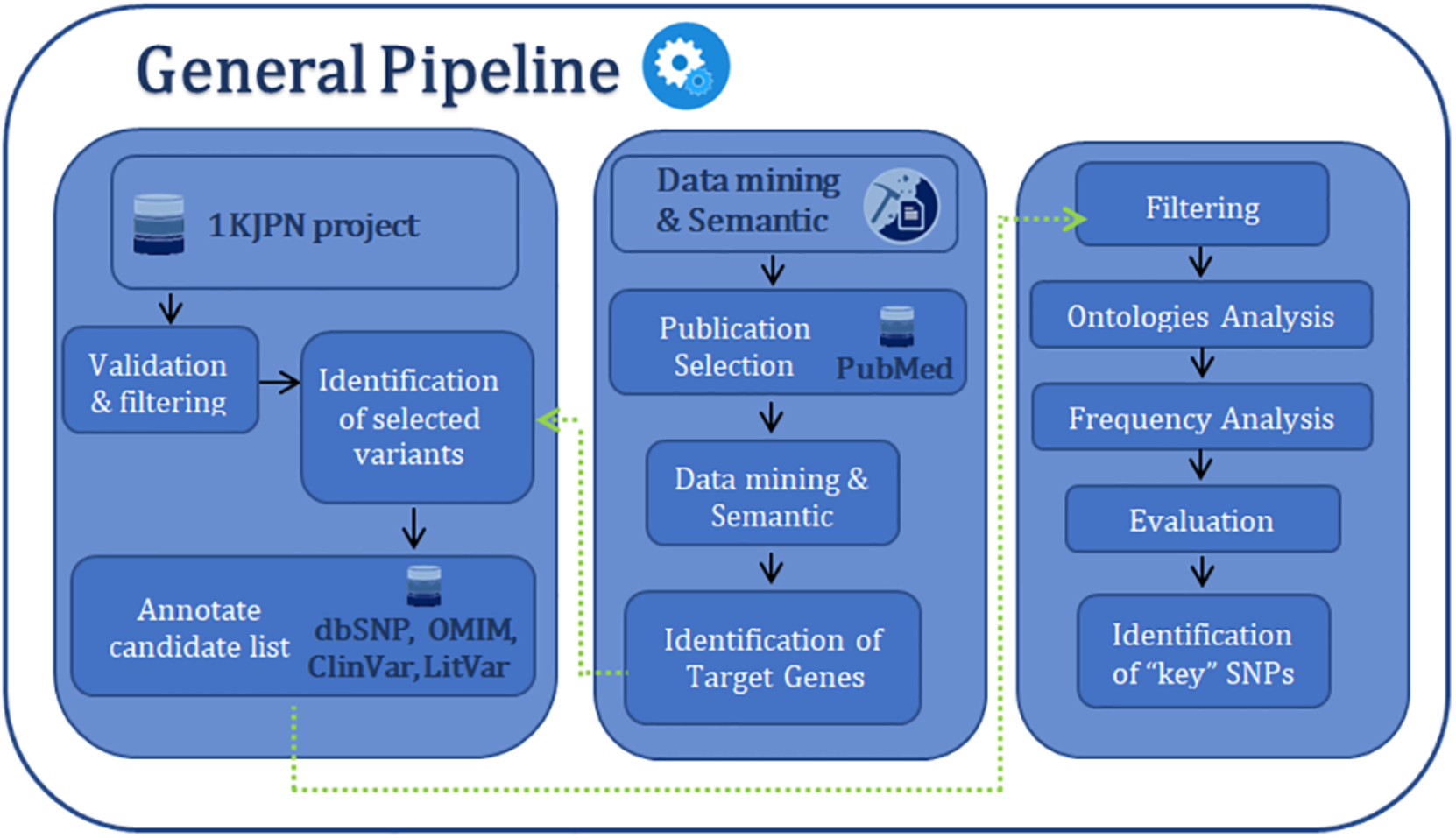
The main procedure pipeline.

## Materials and Methods

### The Dataset

The dataset used was the 2017 update of the 1KJPN project (28), and featured the fully sequenced exome of 3,554 Japanese individuals. The dataset received had already undergone a filtering procedure (Table 1), with the (Single Nucleotide Polymorphisms) SNPs used having ‘passed’ every filtering step. The current dataset only featured autosomes, therefore factors that were located in sex chromosomes were not present. These SNPs included reference SNP ID based on the dbSNP database if applicable. The genomic position of each SNP was based on the GRCh37/hg19 assembly.

**Table 1.** The filtering steps performed in the 3.5K dataset.

| Category | Total SNVs | Matched SNVs | Description |
| --- | --- | --- | --- |
| *Step 1 (Multi-allelic)* | 50.099.977 | 165.439 | Multi-allelic SNVs in 3.5KJPN but biallelic in 1KJPN and 2KJPN |
| *Step 2* | 49.934.538 | 1.373.119 | Depth filter (in naive call, an alternative variant is detected but disappeared with the sequence depth filter, e.g. miscall with CNV, somatic call or misalignment) |
| *Step 3* | 48.561.419 | 2.835.609 | Depth filter (more than 10% of individuals do not fit into the reliable sequence depth range) |
| *Step 4* | 45.725.810 | 6.969.032 | SNVs in highly repetitive regions |
| *Step 5* | 38.756.778 | 1.267.757 | SNVs that are not detected in other alignment tools and variant callers |
| *Step 6* | 37.489.021 | 421.306 | The SNVs's hardy weinberg equilibrium is less than or equal to 0.00001 |
| *Step 7* | 37.489.021 | 13.032.262 |  |

### Identification of target genes

A data mining and semantic study was performed using GR related publications, and a comprehensive list that features 149 autosomal genes which have essential role in GR function or are prime examples of GRE-containing genes was composed (Supplementary Table 1). These genes contain, among others, nuclear receptors, molecular epigenetic regulators, GR cofactors, and several enzymes. The genomic location of each gene was described based on the GRCh37/hg19 assembly. The GeneMANIA webtool (29) was then used towards to estimating the internal links, the biological functions and the biological pathways among genes of interest.

### SNPs Filtering, Annotation and Analysis

Each gene genomic region was then pinpointed in the dataset, and all relative SNPs were extracted. A sliding window algorithm was then used to obtain SNPs that have a reference SNP ID number and are present in the dbSNP database (30). The found SNPs’ genomic location was then updated to be in accordance with the current GRCh38.p13 assembly. All the extracted SNPs were stored in a structured database with all relevant information from the primary dataset including gene name, genetic position, change, and frequency of occurrence based on the sample.

All the identified SNPs were then annotated with relative information from the dbSNPs database (31), LitVar (32), OMIM (33) and ClinVar database (34) (Figure 4). Several types of information were extracted and included in the database produced, using a set of rules based on each database protocols. Particularly, ClinVAR database was used to find possible associations with human health, LitVar database to find the most co-occurred entities regarding diseases, chemicals, and variants, OMIM to extract the genetic disorders based on the corresponding gene and the dbSNP database to find the SNP’s location (intros /exons), common changes, and the allele frequency in different populations. This study’s major goal was to retrieve all the necessary information towards understanding the molecular mechanisms of the stress response system and the SNP in question. The above results were then used to compare the characteristics of the specific dataset to other populations with the help of dbSNP. A second level of filtering analysis then has been performed towards to extracting all the SNPs that are contained in the ClinVar database. Afterwards, based on the results and the available information from the annotation process an ontologies analysis has been performed and the SNPs have been evaluated based on the ClinVar database information (Figure 3). Finally, summarizing all the information collected for each SNPs, a comparison with a dataset featuring Korean individuals was conducted in order to identify attributes specific to the Japanese population that are associated with the GR interactome.

**Figure 4.**
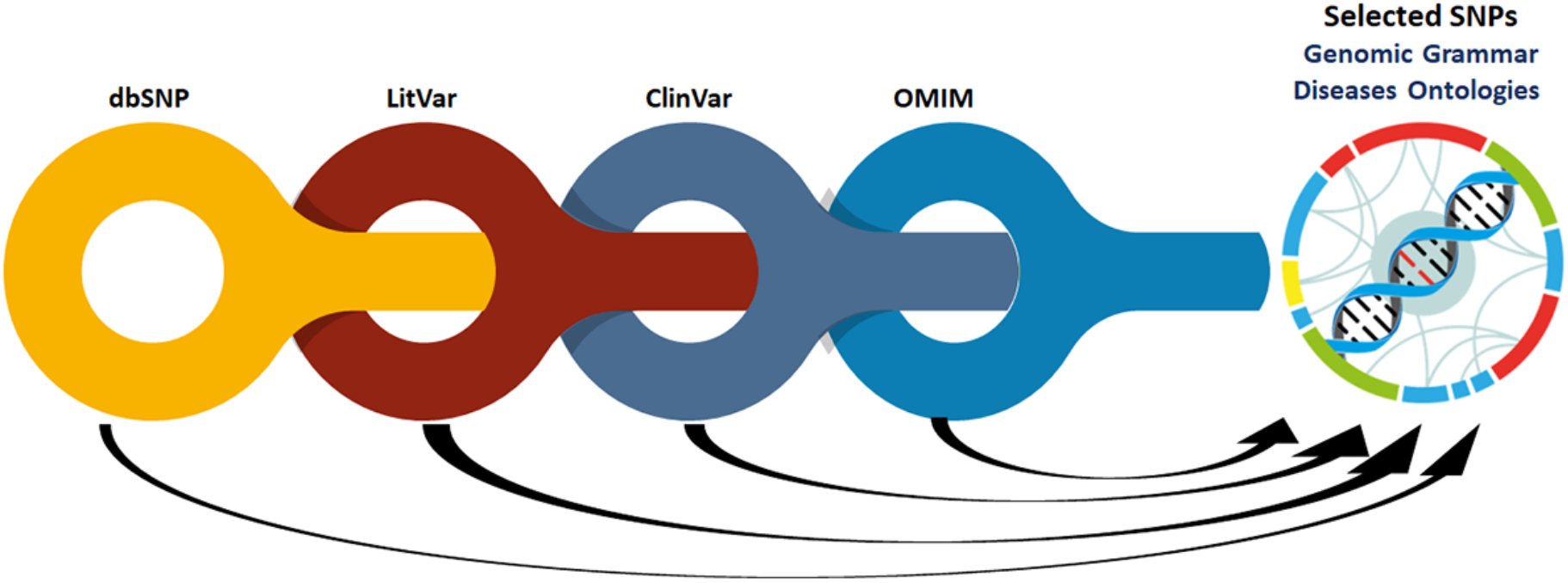
Data Mining and Semantic of selected SNPs.

## Results

The genes checked amounted to 31600 SNPs with a known rs ID that were present in the dbSNP database. Out of 31600 SNPs, 411 were present in the ClinVar database and were chosen as possible SNPs of interest. An ontology analysis based on the corresponding LitVar entries was conducted on mentioned SNPs in order to paint a general picture of the mostly studied mechanisms when it comes to the GR interactome (Figure 5). The results showcase a study focus on metabolic disorders, various neoplasms, and psychiatric disorders. Out of the above 411 polymorphisms, 46 SNPs showcased a possible association with human health or disease according to the ClinVAR database (Table 2). These SNPs were associated with drug metabolism and metabolic disorders, something not surprising since GR has an essential role in metabolism and seems to influence cytochrome P450 function (35, 36). It should also be mentioned that several of the drugs whose metabolism is altered by mentioned SNPs are antidepressants, an observation that is in accordance with GR’s role in neuropsychiatric disorders, and more specifically, depression (37). Lastly, an association with diseases such as chronic obstructive pulmonary disease and inflammatory bowel disease was present in the SNPs, which are characteristic inflammatory diseases, where glucocorticoids are used as potent anti-inflammatory medication (38). These SNPs were then checked on the LitVar database in an effort to exert more information regarding their role in glucocorticoid receptor signaling and homeostasis (Supplementary Table 2).

**Figure 5.**
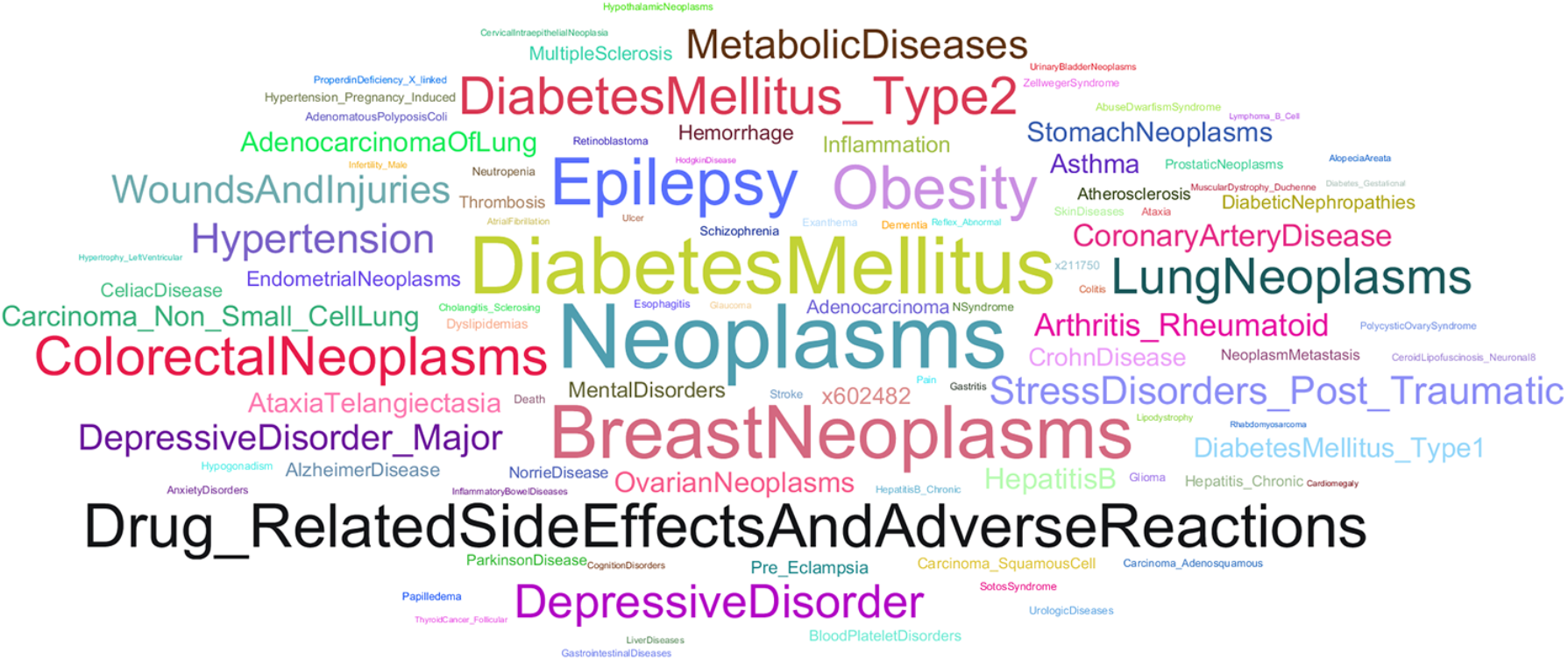
Word Cloud visual representation of the frequency of the disease terms within the final dataset of SNPs associated with glucocorticoid receptor interactome.\

**Table 2.**
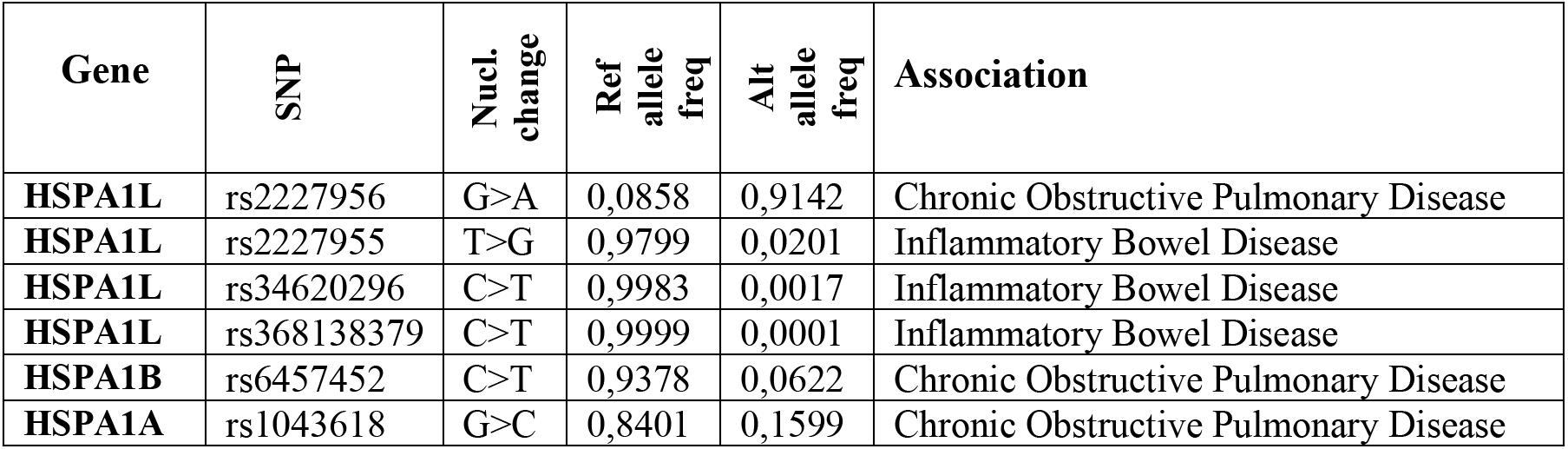
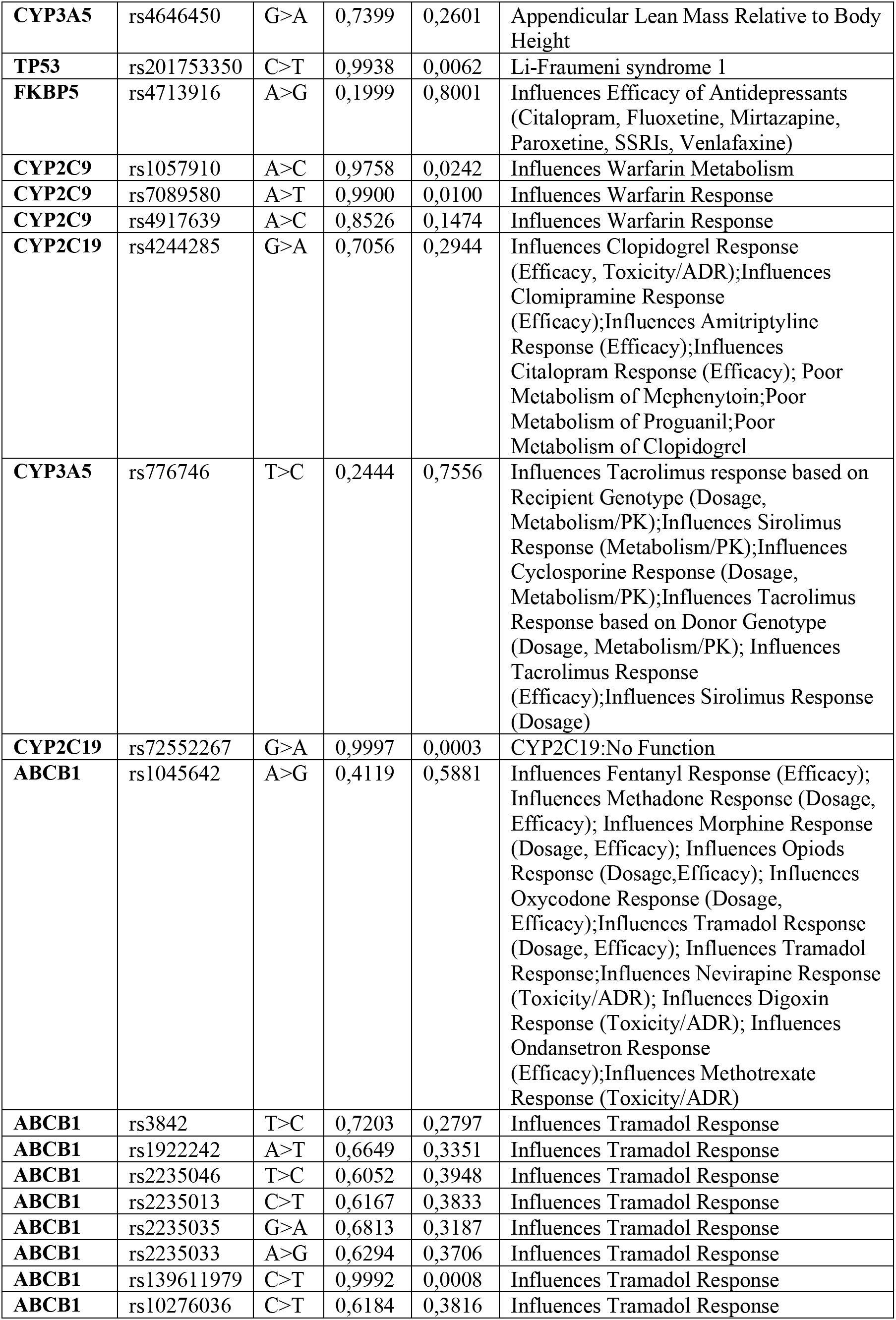
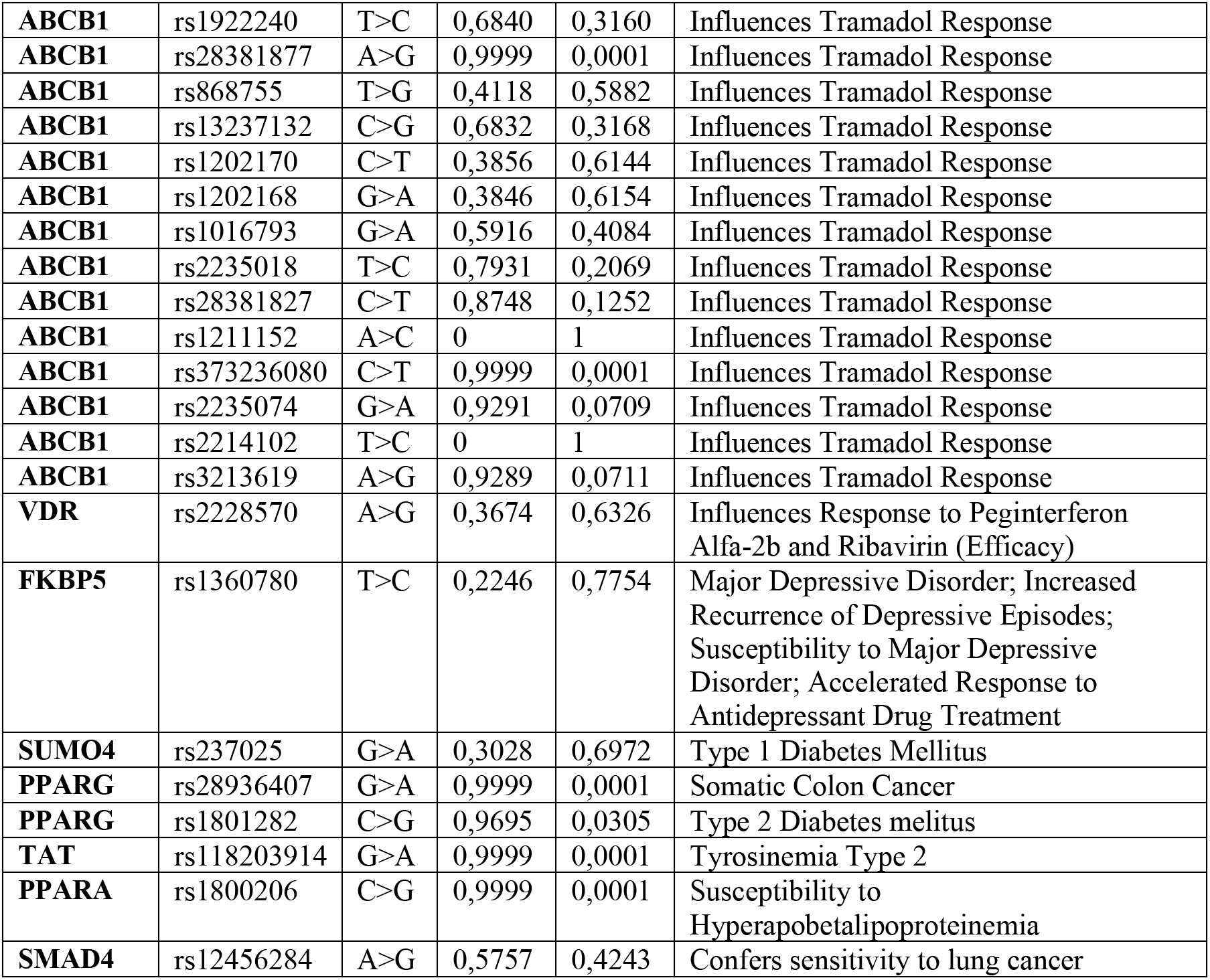
SNPs with a potential pathological association in the ClinVar database, their frequency in the 3.5K dataset, and their association with pathological conditions or physiological mechanisms as stated in the ClinVar database.

| Gene | SNP | Nucl. change | Ref allele freq | Alt allele freq | Association |
| --- | --- | --- | --- | --- | --- |
| HSPA1L | rs2227956 | G>A | 0,0858 | 0,9142 | Chronic Obstructive Pulmonary Disease |
| HSPA1L | rs2227955 | T>G | 0,9799 | 0,0201 | Inflammatory Bowel Disease |
| HSPA1L | rs34620296 | C>T | 0,9983 | 0,0017 | Inflammatory Bowel Disease |
| HSPA1L | rs368138379 | C>T | 0,9999 | 0,0001 | Inflammatory Bowel Disease |
| HSPA1B | rs6457452 | C>T | 0,9378 | 0,0622 | Chronic Obstructive Pulmonary Disease |
| HSPA1A | rs1043618 | G>C | 0,8401 | 0,1599 | Chronic Obstructive Pulmonary Disease |
| **CYP3A5** | rs4646450 | G>A | 0,7399 | 0,2601 | Appendicular Lean Mass Relative to Body Height |
| **TP53** | rs201753350 | C>T | 0,9938 | 0,0062 | Li-Fraumeni syndrome 1 |
| **FKBP5** | rs4713916 | A>G | 0,1999 | 0,8001 | Influences Efficacy of Antidepressants (Citalopram, Fluoxetine, Mirtazapine, Paroxetine, SSRIs, Venlafaxine) |
| **CYP2C9** | rs1057910 | A>C | 0,9758 | 0,0242 | Influences Warfarin Metabolism |
| **CYP2C9** | rs7089580 | A>T | 0,9900 | 0,0100 | Influences Warfarin Response |
| **CYP2C9** | rs4917639 | A>C | 0,8526 | 0,1474 | Influences Warfarin Response |
| **CYP2C19** | rs4244285 | G>A | 0,7056 | 0,2944 | Influences Clopidogrel Response (Efficacy, Toxicity/ADR);Influences Clomipramine Response (Efficacy);Influences Amitriptyline Response (Efficacy);Influences Citalopram Response (Efficacy); Poor Metabolism of Mephenytoin;Poor Metabolism of Proguanil;Poor Metabolism of Clopidogrel |
| **CYP3A5** | rs776746 | T>C | 0,2444 | 0,7556 | Influences Tacrolimus response based on Recipient Genotype (Dosage, Metabolism/PK);Influences Sirolimus Response (Metabolism/PK);Influences Cyclosporine Response (Dosage, Metabolism/PK);Influences Tacrolimus Response based on Donor Genotype (Dosage, Metabolism/PK); Influences Tacrolimus Response (Efficacy);Influences Sirolimus Response (Dosage) |
| **CYP2C19** | rs72552267 | G>A | 0,9997 | 0,0003 | CYP2C19:No Function |
| **ABCB1** | rs1045642 | A>G | 0,4119 | 0,5881 | Influences Fentanyl Response (Efficacy); Influences Methadone Response (Dosage, Efficacy); Influences Morphine Response (Dosage, Efficacy); Influences Opioids Response (Dosage,Efficacy); Influences Oxycodone Response (Dosage, Efficacy);Influences Tramadol Response (Dosage, Efficacy); Influences Tramadol Response;Influences Nevirapine Response (Toxicity/ADR); Influences Digoxin Response (Toxicity/ADR); Influences Ondansetron Response (Efficacy);Influences Methotrexate Response (Toxicity/ADR) |
| **ABCB1** | rs3842 | T>C | 0,7203 | 0,2797 | Influences Tramadol Response |
| **ABCB1** | rs1922242 | A>T | 0,6649 | 0,3351 | Influences Tramadol Response |
| **ABCB1** | rs2235046 | T>C | 0,6052 | 0,3948 | Influences Tramadol Response |
| **ABCB1** | rs2235013 | C>T | 0,6167 | 0,3833 | Influences Tramadol Response |
| **ABCB1** | rs2235035 | G>A | 0,6813 | 0,3187 | Influences Tramadol Response |
| **ABCB1** | rs2235033 | A>G | 0,6294 | 0,3706 | Influences Tramadol Response |
| **ABCB1** | rs139611979 | C>T | 0,9992 | 0,0008 | Influences Tramadol Response |
| **ABCB1** | rs10276036 | C>T | 0,6184 | 0,3816 | Influences Tramadol Response |
| **ABCB1** | rs1922240 | T>C | 0,6840 | 0,3160 | Influences Tramadol Response |
| **ABCB1** | rs28381877 | A>G | 0,9999 | 0,0001 | Influences Tramadol Response |
| **ABCB1** | rs868755 | T>G | 0,4118 | 0,5882 | Influences Tramadol Response |
| **ABCB1** | rs13237132 | C>G | 0,6832 | 0,3168 | Influences Tramadol Response |
| **ABCB1** | rs1202170 | C>T | 0,3856 | 0,6144 | Influences Tramadol Response |
| **ABCB1** | rs1202168 | G>A | 0,3846 | 0,6154 | Influences Tramadol Response |
| **ABCB1** | rs1016793 | G>A | 0,5916 | 0,4084 | Influences Tramadol Response |
| **ABCB1** | rs2235018 | T>C | 0,7931 | 0,2069 | Influences Tramadol Response |
| **ABCB1** | rs28381827 | C>T | 0,8748 | 0,1252 | Influences Tramadol Response |
| **ABCB1** | rs1211152 | A>C | 0 | 1 | Influences Tramadol Response |
| **ABCB1** | rs373236080 | C>T | 0,9999 | 0,0001 | Influences Tramadol Response |
| **ABCB1** | rs2235074 | G>A | 0,9291 | 0,0709 | Influences Tramadol Response |
| **ABCB1** | rs2214102 | T>C | 0 | 1 | Influences Tramadol Response |
| **ABCB1** | rs3213619 | A>G | 0,9289 | 0,0711 | Influences Tramadol Response |
| **VDR** | rs2228570 | A>G | 0,3674 | 0,6326 | Influences Response to Peginterferon Alfa-2b and Ribavirin (Efficacy) |
| **FKBP5** | rs1360780 | T>C | 0,2246 | 0,7754 | Major Depressive Disorder; Increased Recurrence of Depressive Episodes; Susceptibility to Major Depressive Disorder; Accelerated Response to Antidepressant Drug Treatment |
| **SUMO4** | rs237025 | G>A | 0,3028 | 0,6972 | Type 1 Diabetes Mellitus |
| **PPARG** | rs28936407 | G>A | 0,9999 | 0,0001 | Somatic Colon Cancer |
| **PPARG** | rs1801282 | C>G | 0,9695 | 0,0305 | Type 2 Diabetes mellitus |
| **TAT** | rs118203914 | G>A | 0,9999 | 0,0001 | Tyrosinemia Type 2 |
| **PPARA** | rs1800206 | C>G | 0,9999 | 0,0001 | Susceptibility to Hyperapobetalipoproteinemia |
| **SMAD4** | rs12456284 | A>G | 0,5757 | 0,4243 | Confers sensitivity to lung cancer |

Four ABCB1 variations out of the 46 ClinVar entries selected did not also display a LitVar entry. Those are rs373236080, rs28381827, rs28381877, and rs139611979. This discrepancy may be due to the fact that ClinVar also integrates information beyond literature-described associations, such as data from clinical testing labs (39). The resulting SNPs which showcase entries with pathological associations in both the ClinVar and LitVar database are termed SNPs of interest. The results are in accordance with the information received by ClinVar. Some novel associations with various neoplasms emerge, though those are mostly limited to the ABCB1 SNPs, with mentioned gene coding for the P-glycoprotein, whose role in cancer multidrug resistance has been extensively studied in the scientific literature (40).

The frequencies of SNPs of interest which are present in the LitVar database were then characterized based on the nucleotide change region and type of mutation and later compared with a dataset consisting of 1465 Korean individuals, since those two populations display somewhat high similarity (Table 3), and possible differences may display distinct genetic characteristics that may influence GR-associated or stress-associated processes (41). Most of the SNPs compared are located in intronic regions. Although introns were thought to be of small biological importance, modern studies have shown that they have a great role in essential processes from alternate splicing to regulating gene expression (42). The comparison among the Japanese and Korean populations highlighted the rs1043618 as a polymorphism with a substantially different frequency among populations. This polymorphism has been associated with COPD in ClinVar and Depression in LitVar.

**Table 3.** A genetic comparison between the Japanese and Korean population focusing on SNPs of interest frequency.

| Gene | SNP | Nucleotide change | Nucleotide change region | Nucleotide frequency in Japanese population | Nucleotide frequency in Korean population |
| --- | --- | --- | --- | --- | --- |
| HSPA1L | rs2227956 | G>A | Missense variant | G=0,0858 | G=0,0765 |
| HSPA1L | rs2227955 | T>G | Missense variant | G=0,0201 | G=0,0171 |
| HSPA1L | rs34620296 | C>T | Missense variant | T=0,0017 | T= 0,0048 |
| HSPA1L | rs368138379 | C>T | Missense variant | T=0,0001 | - |
| HSPA1B | rs6457452 | C>T | 5 Prime UTR Variant | T=0,0622 | T=0,0875 |
| HSPA1A | rs1043618 | G>C | 5 Prime UTR Variant | C=0,1599 | C=0,2801 |
| CYP3A5 | rs4646450 | G>A | Intron Variant | A=0,2601 | A=0,2304 |
| TP53 | rs201753350 | C>T | Missense Variant | T=0,0062 | T=0,0055 |
| FKBP5 | rs4713916 | A>G | Intron Variant | A=0,1999 | A=0,2096 |
| CYP2C9 | rs1057910 | A>C | Missense Variant | C=0,0242 | C=0,0413 |
| CYP2C9 | rs7089580 | A>T | Intron Variant | T=0,01 | T=0,0082 |
| CYP2C9 | rs4917639 | A>C | Intron Variant | C=0, 1474 | C=0,1345 |
| CYP2C19 | rs4244285 | G>A | Synonymous Variant | A=0,2944 | A=0,2765 |
| CYP3A5 | rs776746 | T>C | Splice Acceptor Variant | C=0,2444 | C=0,2249 (1K) |
| CYP2C19 | rs72552267 | G>A | Missense Variant | A=0,0003 | - |
| ABCB1 | rs1045642 | A>G | Missense Variant | A=0,4119 | A=0,3488 |
| ABCB1 | rs3842 | T>C | 3 Prime UTR Variant | C=0,2797 | C=0,3061 |
| ABCB1 | rs1922242 | A>T | Intron Variant | T=0,3351 | T=0,3717 |
| ABCB1 | rs2235046 | T>C | Intron Variant | C=0,3948 | C=0,4085 |
| ABCB1 | rs2235013 | C>T | Intron Variant | T=0,3833 | T=0,4065 |
| ABCB1 | rs2235035 | G>A | Intron Variant | A=0,3187 | A=0,3590 |
| ABCB1 | rs2235033 | A>G | Intron Variant | G=0, 3706 | G=0,4065 |
| ABCB1 | rs10276036 | C>T | Intron Variant | T=0,3816 | T=0,4061 |
| ABCB1 | rs1922240 | T>C | Intron Variant | C=0,3160 | C=0,3573 |
| ABCB1 | rs868755 | T>G | Intron Variant | T=0,4118 | T=0,3788 |
| ABCB1 | rs13237132 | C>G | Intron Variant | G=0,3168 | G=0,3563 |
| ABCB1 | rs1202170 | C>T | Intron Variant | C=0,3856 | C=0,4058 |
| ABCB1 | rs1202168 | G>A | Intron Variant | G=0,3846 | G=0,4038 |
| ABCB1 | rs1016793 | G>A | Intron Variant | A=0,4084 | A=0,3860 |
| ABCB1 | rs2235018 | T>C | Intron Variant | C=0,2069 | C=0,2160 |
| ABCB1 | rs1211152 | A>C | Intron Variant | A=0 | A=0,001 |
| ABCB1 | rs2235074 | G>A | Intron Variant | A=0,0709 | A=0,0565 |
| ABCB1 | rs2214102 | T>C | Synonymous Variant | T=0 | T=0,0003 |
| ABCB1 | rs3213619 | A>G | Intron Variant | G=0,0709 | G=0,0561 |
| VDR | rs2228570 | A>G | Initiator Codon Variant | A=0,3674 | A=0,4041 |
| FKBP5 | rs1360780 | T>C | Intron Variant | T=0,2246 | T=0,2392 |
| SUMO4 | rs237025 | G>A | Missense Variant | G=0,3028 | G=0,2973 |
| PPARG | rs28936407 | G>A | Missense Variant | A=0,0001 | - |
| PPARG | rs1801282 | C>G | Missense Variant | G=0,0305 | G=0,0517 |
| TAT | rs118203914 | G>A | Stop Gained | A=0,0001 | - |
| PPARA | rs1800206 | C>G | Missense Variant | G=0,0001 | G=0,0005 (1K) |
| SMAD4 | rs12456284 | A>G | 3 Prime UTR Variant | 0,4243 | G=0,4049 |

## Discussion

The current study restates the importance of the stress response system in human pathophysiology. Polymorphisms on genes characteristic of the GR interactome lead to metabolic, psychiatric, cancer and inflammatory diseases. These results are in accordance with the stress response system’s role in neuropsychiatric disorders (43, 44) and the important role of the glucocorticoid receptor in inflammation (45) and metabolism(46). A peculiar observation was that according to LitVar several SNPs were the focus of multiple cancer studies, though ClinVar pathological associations with cancer were minimal. This observation may be due to several factors, such as the scientific community’s focus on cancer research or a possibly currently emerging association between the GR interactome and various neoplasms. The importance of introns in several pathophysiological conditions was also highlighted, since the vast majority of found SNPs were located in intronic regions. Moreover, the similarities in SNP frequencies between the Korean and Japanese populations are in accordance with the fact that mainland Japanese are genetically close to Koreans (47). Nonetheless, an interesting discrepancy was present between these two populations, which extended to discrepancies with the frequencies present on the TOPMED and 1000 Genomes Project. Japanese individuals showcased a rs1043618 frequency of 0,1599 while Koreans had a frequency of 0,2801, with the TOPMED and 1000 Genomes Project frequencies being 0,478474 and 0,4812, respectively. This HSPA1A polymorphism may influence Hsp70 protein levels through translation efficiency alterations or post-transcriptional regulation (48). This polymorphism’s potential role in COPD in response to environmental stimuli deserves special mention. Specifically, rs1043618 has been associated with susceptibility to chronic obstructive pulmonary disease (COPD) in response to environmental stressors in a Mexican population (49). This effect may be due to the fact that HSPA1A codes a 70kDa Hsp protein that partakes in the GR chaperone complex, and possible protein level alterations may lead to a problematic response to stressors. This observation is really intriguing since COPD displays a higher incidence rate in Koreans than in Japanese populations while smoking trends between those countries are quite similar (50, 51). This may lead to the speculation that the rs1043618 could be partially responsible for such a phenomenon. Nevertheless, it is important to state that specific SNPs may be associated with a disease in one population but show no association in another (52). All in all, researching the genetic intricacies of the GR interactome in different population may provide new target genes for the management or treatment of stress related pathologies.

## Supporting information

https://www.dropbox.com/s/fbhx95mrcyubps9/DVlachakis_MS_JAPAN_Supplementary.pdf?dl=0

## Acknowledgments

Not applicable.

## References

1. Chrousos GP: Stress and disorders of the stress system. Nature reviews Endocrinology 5: 374–381, 2009.

2. Charmandari E, Tsigos C and Chrousos G: Endocrinology of the stress response. Annual review of physiology 67: 259–284, 2005.

3. Nicolaides NC, Kyratzi E, Lamprokostopoulou A, Chrousos GP and Charmandari E: Stress, the stress system and the role of glucocorticoids. Neuroimmunomodulation 22: 6–19, 2015.

4. Chrousos GP: Stressors, stress, and neuroendocrine integration of the adaptive response. The 1997 Hans Selye Memorial Lecture. Annals of the New York Academy of Sciences 851: 311–335, 1998.

5. Ramamoorthy S and Cidlowski JA: Corticosteroids: Mechanisms of Action in Health and Disease. Rheum Dis Clin North Am 42: 15–vii, 2016.

6. Papageorgiou L, Shalzi L, Pierouli K, Papakonstantinou E, Manias S, Dragoumani K, Nicolaides NC, Giannakakis A, Bacopoulou F, et al.: An updated evolutionary study of the nuclear receptor protein family. World Acad Sci J 3: 51, 2021.

7. Papageorgiou L, Shalzi L, Efthimiadou A, Bacopoulou F, Chrousos GP, Eliopoulos E and Vlachakis D: Conserved functional motifs of the nuclear receptor superfamily as potential pharmacological targets. Int J Epigen 1: 3, 2021.

8. Stephens MA and Wand G: Stress and the HPA axis: role of glucocorticoids in alcohol dependence. Alcohol research: current reviews 34: 468–483, 2012.

9. Mitsis T, Papageorgiou L, Efthimiadou A, Bacopoulou F, Vlachakis D, Chrousos GP, Eliopoulos E, Mitsis T, Papageorgiou L, et al.: A comprehensive structural and functional analysis of the ligand binding domain of the nuclear receptor superfamily reveals highly conserved signaling motifs and two distinct canonical forms through evolution. World Acad Sci J 1: 264–274, 2019.

10. Thau L, Gandhi J and Sharma S: Physiology, Cortisol. In: StatPearls. StatPearls Publishing Copyright © 2021, StatPearls Publishing LLC., Treasure Island (FL), 2021.

11. Benarroch EE: The locus ceruleus norepinephrine system: functional organization and potential clinical significance. Neurology 73: 1699–1704, 2009.

12. Lü Y-F, Yang Y, Li C-L, Wang Y, Li Z and Chen J: The Locus Coeruleus-Norepinephrine System Mediates Empathy for Pain through Selective Up-Regulation of P2X3 Receptor in Dorsal Root Ganglia in Rats. Front Neural Circuits 11: 66–66, 2017.

13. Kanczkowski W, Sue M and Bornstein SR: Adrenal Gland Microenvironment and Its Involvement in the Regulation of Stress-Induced Hormone Secretion during Sepsis. Frontiers in endocrinology 7: 156, 2016.

14. McEwen BS, Bowles NP, Gray JD, Hill MN, Hunter RG, Karatsoreos IN and Nasca C: Mechanisms of stress in the brain. Nat Neurosci 18: 1353–1363, 2015.

15. Scheschowitsch K, Leite JA and Assreuy J: New Insights in Glucocorticoid Receptor Signaling-More Than Just a Ligand-Binding Receptor. Frontiers in endocrinology 8: 16–16, 2017.

16. Sevilla LM and Pérez P: Roles of the Glucocorticoid and Mineralocorticoid Receptors in Skin Pathophysiology. Int J Mol Sci 19: 1906, 2018.

17. Koning A-SCAM, Buurstede JC, van Weert LTCM and Meijer OC: Glucocorticoid and Mineralocorticoid Receptors in the Brain: A Transcriptional Perspective. J Endocr Soc 3: 1917–1930, 2019.

18. Sever R and Glass CK: Signaling by nuclear receptors. Cold Spring Harb Perspect Biol 5: a016709–a016709, 2013.

19. Mazaira GI, Zgajnar NR, Lotufo CM, Daneri-Becerra C, Sivils JC, Soto OB, Cox MB and Galigniana MD: The Nuclear Receptor Field: A Historical Overview and Future Challenges. Nucl Receptor Res 5: 101320, 2018.

20. Heitzer MD, Wolf IM, Sanchez ER, Witchel SF and DeFranco DB: Glucocorticoid receptor physiology. Reviews in endocrine & metabolic disorders 8: 321–330, 2007.

21. Phuc Le P, Friedman JR, Schug J, Brestelli JE, Parker JB, Bochkis IM and Kaestner KH: Glucocorticoid receptor-dependent gene regulatory networks. PLoS Genet 1: e16–e16, 2005.

22. Timmermans S, Souffriau J and Libert C: A General Introduction to Glucocorticoid Biology. Frontiers in Immunology 10: 2019.

23. Frego L and Davidson W: Conformational changes of the glucocorticoid receptor ligand binding domain induced by ligand and cofactor binding, and the location of cofactor binding sites determined by hydrogen/deuterium exchange mass spectrometry. Protein Sci 15: 722–730, 2006.

24. Vandevyver S, Dejager L and Libert C: On the trail of the glucocorticoid receptor: into the nucleus and back. Traffic (Copenhagen, Denmark) 13: 364–374, 2012.

25. Petta I, Dejager L, Ballegeer M, Lievens S, Tavernier J, De Bosscher K and Libert C: The Interactome of the Glucocorticoid Receptor and Its Influence on the Actions of Glucocorticoids in Combatting Inflammatory and Infectious Diseases. Microbiol Mol Biol Rev 80: 495–522, 2016.

26. Petta I, Dejager L, Ballegeer M, Lievens S, Tavernier J, De Bosscher K and Libert C: The Interactome of the Glucocorticoid Receptor and Its Influence on the Actions of Glucocorticoids in Combatting Inflammatory and Infectious Diseases. Microbiology and Molecular Biology Reviews 80: 495–522, 2016.

27. Pooley JR, Rivers CA, Kilcooley MT, Paul SN, Cavga AD, Kershaw YM, Muratcioglu S, Gursoy A, Keskin O, et al.: Beyond the heterodimer model for mineralocorticoid and glucocorticoid receptor interactions in nuclei and at DNA. PLOS ONE 15: e0227520, 2020.

28. Nagasaki M, Yasuda J, Katsuoka F, Nariai N, Kojima K, Kawai Y, Yamaguchi-Kabata Y, Yokozawa J, Danjoh I, et al.: Rare variant discovery by deep whole-genome sequencing of 1,070 Japanese individuals. Nature Communications 6: 8018, 2015.

29. Warde-Farley D, Donaldson SL, Comes O, Zuberi K, Badrawi R, Chao P, Franz M, Grouios C, Kazi F, et al.: The GeneMANIA prediction server: biological network integration for gene prioritization and predicting gene function. Nucleic Acids Research 38: W214–W220, 2010.

30. Sherry ST, Ward MH, Kholodov M, Baker J, Phan L, Smigielski EM and Sirotkin K: dbSNP: the NCBI database of genetic variation. Nucleic acids research 29: 308–311, 2001.

31. Smigielski EM, Sirotkin K, Ward M and Sherry ST: dbSNP: a database of single nucleotide polymorphisms. Nucleic Acids Res 28: 352–355, 2000.

32. Allot A, Peng Y, Wei CH, Lee K, Phan L and Lu Z: LitVar: a semantic search engine for linking genomic variant data in PubMed and PMC. Nucleic Acids Res 46: W530–W536, 2018.

33. Hamosh A, Scott AF, Amberger J, Bocchini C, Valle D and McKusick VA: Online Mendelian Inheritance in Man (OMIM), a knowledgebase of human genes and genetic disorders. Nucleic Acids Res 30: 52–55, 2002.

34. Landrum MJ, Chitipiralla S, Brown GR, Chen C, Gu B, Hart J, Hoffman D, Jang W, Kaur K, et al.: ClinVar: improvements to accessing data. Nucleic Acids Res 48: D835D844, 2020.

35. Dvorak Z and Pavek P: Regulation of drug-metabolizing cytochrome P450 enzymes by glucocorticoids. Drug metabolism reviews 42: 621–635, 2010.

36. Rose AJ, Vegiopoulos A and Herzig S: Role of glucocorticoids and the glucocorticoid receptor in metabolism: insights from genetic manipulations. J Steroid Biochem Mol Biol 122: 10–20, 2010.

37. Kadmiel M and Cidlowski JA: Glucocorticoid receptor signaling in health and disease. Trends Pharmacol Sci 34: 518–530, 2013.

38. Coutinho AE and Chapman KE: The anti-inflammatory and immunosuppressive effects of glucocorticoids, recent developments and mechanistic insights. Molecular and cellular endocrinology 335: 2–13, 2011.

39. Henrie A, Hemphill SE, Ruiz-Schultz N, Cushman B, DiStefano MT, Azzariti D, Harrison SM, Rehm HL and Eilbeck K: ClinVar Miner: Demonstrating utility of a Webbased tool for viewing and filtering ClinVar data. Hum Mutat 39: 1051–1060, 2018.

40. Robinson K and Tiriveedhi V: Perplexing Role of P-Glycoprotein in Tumor Microenvironment. Front Oncol 10: 265–265, 2020.

41. Takeuchi F, Katsuya T, Kimura R, Nabika T, Isomura M, Ohkubo T, Tabara Y, Yamamoto K, Yokota M, et al.: The fine-scale genetic structure and evolution of the Japanese population. PLoS One 12: e0185487–e0185487, 2017.

42. Jo B-S and Choi SS: Introns: The Functional Benefits of Introns in Genomes. Genomics Inform 13: 112–118, 2015.

43. Jacobson L: Hypothalamic-pituitary-adrenocortical axis: neuropsychiatric aspects. Comprehensive Physiology 4: 715–738, 2014.

44. Godoy LD, Rossignoli MT, Delfino-Pereira P, Garcia-Cairasco N and de Lima Umeoka EH: A Comprehensive Overview on Stress Neurobiology: Basic Concepts and Clinical Implications. Frontiers in behavioral neuroscience 12: 127, 2018.

45. Hübner S, Dejager L, Libert C and Tuckermann JP: The glucocorticoid receptor in inflammatory processes: transrepression is not enough. Biological chemistry 396: 1223–1231, 2015.

46. Akalestou E, Genser L and Rutter GA: Glucocorticoid Metabolism in Obesity and Following Weight Loss. Frontiers in Endocrinology 11: 2020.

47. Watanabe Y, Naka I, Khor S-S, Sawai H, Hitomi Y, Tokunaga K and Ohashi J: Analysis of whole Y-chromosome sequences reveals the Japanese population history in the Jomon period. Scientific Reports 9: 8556, 2019.

48. He M, Guo H, Yang X, Zhang X, Zhou L, Cheng L, Zeng H, Hu FB, Tanguay RM, et al.: Functional SNPs in HSPA1A gene predict risk of coronary heart disease. PloS one 4: e4851–e4851, 2009.

49. Ambrocio-Ortiz E, Pérez-Rubio G, Ramírez-Venegas A, Hernández-Zenteno R, Del Angel-Pablo AD, Pérez-Rodríguez ME, Salazar AM, Abarca-Rojano E and Falfán-Valencia R: Effect of SNPs in HSP Family Genes, Variation in the mRNA and Intracellular Hsp Levels in COPD Secondary to Tobacco Smoking and Biomass-Burning Smoke. Frontiers in Genetics 10: 2020.

50. Leem AY, Park B, Kim YS, Jung JY and Won S: Incidence and risk of chronic obstructive pulmonary disease in a Korean community-based cohort. International journal of chronic obstructive pulmonary disease 13: 509–517, 2018.

51. Funatogawa I, Funatogawa T and Yano E: Trends in smoking and lung cancer mortality in Japan, by birth cohort, 1949-2010. Bull World Health Organ 91: 332–340, 2013.

52. Rao S, Yao Y, Ryan J, Li T, Wang D, Zheng C, Xu Y and Xu Q: Common variants in FKBP5 gene and major depressive disorder (MDD) susceptibility: a comprehensive meta-analysis. Sci Rep 6: 32687–32687, 2016.

