## Supplementary material for "A genomic study of the Japanese population focusing on the glucocorticoid receptor interactome highlights distinct genetic characteristics associated with stress response": https://www.dropbox.com/s/fbhx95mrcyubps9/DVlachakis_MS_JAPAN_Supplementary.pdf?dl=0

**Supplementary Table 1.** A literature-based list of GR interacting autosomal genes. The table features the gene in question, its influence on GR function or vice versa, the Pubmed ID of the paper pertaining to mentioned influence, and the gene's location in the human genome.

| Factor | Influence | Pubmed ID | Chromosome and Position |
|---|---|---|---|
| **BAG1** | Interacts with hsp70, binds to hinge region, inhibits DNA binding and transactivation | 9603979<br>11101523 | 9 NC_000009.11 (33252469..33264759, complement) |
| **CDKs (1,2,5)** | Different effects on GRE-containing promoters, based on phosphorylation sites (Serine 203 and Serine 226 decreased activation, Serine 211 enhanced activation) | 19787703 | 10 NC_000010.10 (62538089..62554610)<br><br>12 NC_000012.11 (56360556..56366573)<br><br>7 NC_000007.13 (150750899..150755052, complement) |
| **TAT** | GR is essential for TAT gene induction. | 11420718 | 16 NC_000016.9 (71600754..71610998, complement) |
| **GSK-3b** | Leads to a conformation change in GR, attenuates GC signaling | 19787703 | 3 NC_000003.11 (119540800..119813264, complement) |
| **ERK2** | Decreases receptor activity | 9199329 | 22 NC_000022.10 (22113946..22221970, complement) |
| **p38 (MAPK11-14)** | Enhances GRE-related activity | 15817653 | 22 NC_000022.10 (50702142..50708779, complement)<br><br>22 NC_000022.10 (50691331..50700248, complement)<br><br>6 NC_000006.11 (36098261..36112301)<br><br>6 NC_000006.11 (35995412..36079013) |
| **MAPK8(JNK1)** | Decreases receptor activity | 12351702 | 10 NC_000010.10 (49514682..49647403) |
| **Ubc9** | Sumoylation/ Increases GR activity | 12144530 | 16 NC_000016.9 (1357420..1377019) |
| **LCK** | Unliganded GR is part of a TcR-linked multiprotein | 27169854 | 1 NC_000001.10 (32716840..32751766) |

| | | | |
|---|---|---|---|
| | complex containing Hsp90, LCK, and FYN | | |
| **FYN** | Unliganded GR is part of a TcR-linked multiprotein complex containing Hsp90, LCK, and FYN | 27169854 | 6 NC_000006.11 (111981535..112194655, complement) |
| **SUMO1** | Sumoylation/Increased GR activity | 12144530 | 2 NC_000002.11 (203070903..203103322, complement) |
| **CHIP(STUB1)** | Receptor downregulation and decreased transactivation | 15761032 | 16 NC_000016.9 (730115..732768) |
| **Mdm2** | Takes part in GR degradation | 12897156 | 12 NC_000012.11 (69201952..69239324) |
| **DNA-PKcs** | Phosphorylation of GR hinge region | 9038175 | 8 NC_000008.10 (48685669..48872743, complement) |
| **CEBPB** | GR potentiates the action of CEBPB, Along with p21 it can inhibit cdk2 action | 9817600 | 20 NC_000020.10 (48807120..48809227) |
| **HDAC2** | Influences GC sensitivity (overexpression leads to increased sensitivity) | 23953592 | 6 NC_000006.11 (114257320..114292359, complement) |
| **TSC22D3** | GR upregulates its' specific gene | 23953592 | X NC_000023.10 (106956451..107020559, complement) |
| **SGK1** | GR upregulates its' specific gene | 23953592 | 6 NC_000006.11 (134490384..134639196, complement) |
| **VDR** | Features GREs and glucocorticoids regulate its' transcription | 20398752 | 12 NC_000012.11 (48235320..48298814, complement) |
| **ZFP36** | GR upegulates its' specific gene | 23953592 | 19 NC_000019.9 (39897487..39900052) |
| **DUSP1** | GR upegulates its' specific gene | 23953592 | 5 NC_000005.9 (172195093..172198203, complement) |
| **β-arrestin (1,2)** | GR regulates their gene expression (β-arrestin 1 +, β-arrestin 2 -) | 23953592 | 11 NC_000011.9 (74971166..75062875, complement)<br><br>17 NC_000017.10 (4613789..4624795) |
| **BGLAP** | GR downregulates its' specific gene | 23953592 | 1 NC_000001.10 (156211951..156213123) |
| **TBP** | GR's AF-1 domain binds TBP, overexpression of TBP | 16469772 | 6 NC_000006.11 (170863384..170881958) |

| | | | |
|---|---|---|---|
| | leads to stimulated expression of GR-driven reporters | | |
| **CBP** | Coactivator, Interacts with GR and p300 to form docking platform for transcription factors | 19818358 | 16 NC_000016.9 (3775055..3930121, complement) |
| **p300** | Coactivator, Interacts with GR and CBP to form docking platform for transcription factors | 19818358 | 22 NC_000022.10 (41488614..41576081) |
| **Pcaf** | Interacts with p300CBP and STAT3 to stimulate GR activity | 27169854 | 3 NC_000003.11 (20081524..20195896) |
| **NCOAs(1,2,3)** | Coactivators that assist DNA expression, upregulation | 19805480 | 2 NC_000002.11 (24714919..24993571)<br><br>8 NC_000008.10 (71021997..71316062, complement)<br><br>20 NC_000020.10 (46130601..46285621) |
| **SMAD6** | SMAD6 suppresses GR function | 27169854 | 15 NC_000015.9 (66994674..67074338) |
| **DAP3** | Binds HSP90,increases transactivation activity | 10903152 | 1 NC_000001.10 (155657693..155708801) |
| **DAXX** | Suppresses GR expression | 12595526 | 6 NC_000006.11 (33286335..33290793, complement) |
| **PP1 (PPP1CA, PPP1CB, PPP1CC)** | May reverse GR phosphorylation | 19818358 | 11 NC_000011.9 (67165652..67169376, complement)<br><br>2 NC_000002.11 (28974614..29025806)<br><br>12 NC_000012.11 (111157613..111180783, complement) |
| **PP2(PPP2CA, PPP2CB)** | May reverse GR phosphorylation | 19818358 | 5 NC_000005.9 (133532148..133561950, complement)<br><br>8 NC_000008.10 (30643126..30670352, complement) |
| **MED1** | Enhances GR expression | 10508170 | 17 NC_000017.10 (37560538..37607527, complement) |

| | | | |
|---|---|---|---|
| **MED14** | Enhances GR expression | 10508170 | X NC_000023.10 (40508795..40595374, complement) |
| **HNRNPU** | Overexpression of HNRPU leads to GR inactivation | 9353307 | 1 NC_000001.10 (245013602..245027827, complement) |
| **HSP90 (HSP90AA1, HSP90AA2P)** | Essential chaperone for GR function | 28224564 | 14 NC_000014.8 (102547075..102606086, complement)<br><br>11 NC_000011.9 (27909718..27912639, complement) pseudogene |
| **HSP70 (HSPA1A, HSPA1B, HSPA1L)** | Essential chaperone for GR function | 24949977 | 6 NC_000006.11 (31782952..31785719)<br><br>6 NC_000006.11 (31789964..31798032)<br><br>6 NC_000006.11 (31777396..31790093, complement) |
| **HSP40 (DNAJA1, DNAJA2, DNAJA3, DNAJB1)** | Increases the efficiency of the GR/chaperons complex | 24345775<br>24949977<br>20453930<br>26245905 | 9 NC_000009.11 (33025209..33039905)<br><br>16 NC_000016.9 (46989274..47007625, complement)<br><br>16 NC_000016.9 (4475806..4506776)<br><br>19 NC_000019.9 (14625581..14640087, complement) |
| **HOP** | Increases the efficiency of the GR/chaperons complex | 10764743<br>24949977 | 11 NC_000011.9 (63952206..63972020) |
| **p23** | Increases the efficiency of the GR/chaperons complex | 24345775<br>24949977 | 12 NC_000012.11 (57057125..57082138, complement) |
| **MR** | Heterodimerization with GR and coordinates transcription | 11154266 | 4 NC_000004.11 (148999915..149365850, complement) |
| **Cytochrome p450 enzymes (CYP3A4, CYP3A5, CYP2C8,** | GR regulates the enzymes' expression | 24451000 | 7 NC_000007.13 (99354583..99381811, complement) |

| | | | |
|---|---|---|---|
| **CYP2C9, CYP2C19)** | | | 7 NC_000007.13 (99245813..99277636, complement)<br><br>10 NC_000010.10 (96796529..96829255, complement)<br><br>10 NC_000010.10 (96698350..96749486)<br><br>10 NC_000010.10 (96522463..96612671) |
| **P-glycoprotein** | GR regulates its' expression | 24451000 | 7 NC_000007.13 (87133179..87342639, complement) |
| **FKBP4(FKBP52)** | Regulates GR signaling, possibly positive regulation | 19818358 | 12 NC_000012.11 (2904108..2914589) |
| **FKBP5(FKBP51)** | Regulates GR signaling, possibly negative regulation | 19818358 | 6 NC_000006.11 (35541362..35696360, complement) |
| **NRIP1** | Negatively regulates the activity of GR | 12773562 | 21 NC_000021.8 (16333556..16438224, complement) |
| **CLOCK** | Represses GR-induced transcriptional activity | 19818358 | 4 NC_000004.11 (56294068..56413076, complement) |
| **BMAL1** | Represses GR-induced transcriptional activity | 19818358 | 11 NC_000011.9 (13299325..13408813) |
| **AP-1 (specifically c-Fos)** | GR weakly interacts and inhibits AP-1 depended transcription. Specifically, GR binds cFos/cJun via a sequence unique to cFos | 27169854 | 14 NC_000014.8 (75745477..75748937) |
| **NF-κB** | GR interacts with NF-κB through the second zinc finger of the ligand-binding domain and acts negatively on the p65/RelA subunit of NFκB. | 19818358 | 11 NC_000011.9 (65421067..65430443, complement) |
| **POU2F1** | GR interacts with POU2F1 in order to bind to distal nGRE | 9891005 | 1 NC_000001.10 (167190066..167396582) |
| **POU2F2** | GR interacts with POU2F2 in order to promote the binding of POU2F2 o specific sequences | 10480874 | 19 NC_000019.9 (42590262..42636625, complement) |
| **p21** | Along with CEBP, it can inhibit cdk2 action | 11369759 | 6 NC_000006.11 (36644237..36655116) |
| **Smad3** | GR inhibits the transcriptional activation function of Smad3 | 10518526 | 15 NC_000015.9 (67358036..67487533) |

| | | | |
|---|---|---|---|
| **Smad4** | GR inhibits the transcriptional activation function of Smad4 (only in vitro) | 10518526 | 18 NC_000018.9 (48556583..48611412 |
| **RanBP9** | Overexpression of RanBP9 leads to enhanced GR activity | 12361945 | 6 NC_000006.11 (13621730..13711796, complement) |
| **SET** | Acts as ligand-activated GR-responsive transcriptional repressor | 18096310 | 9 NC_000009.11 (131445934..131458675) |
| **NFATc** | GR, through protein-protein interaction, interferes with NFATc ability to bind to specific DNA regions | 10623828 | 18 NC_000018.9 (77155772..77289323) |
| **BAFs (BAF57, BAF60a, BAF250a, BAF250b)** | Human analogs of the SWI/SNF complex. These complexes partake in glucocorticoid stimulated transcription by interacting with GR. | 26278180 | 17 NC_000017.10 (38781214..38805658, complement)<br><br>12 NC_000012.11 (50478760..50494494)<br><br>1 NC_000001.10 (27022522..27108601)<br><br>6 NC_000006.11 (157098980..157531913) |
| **p53** | GR has the ability to inhibit p53-depended functions | 22773829<br>11080152 | 17 NC_000017.10 (7571720..7590868, complement) |
| **PPP5** | Suppression of PP5 results to nuclear accumulation of GR | 11389770 | 19 NC_000019.9 (46850251..46896238) |
| **STAT3** | Acts as transcriptional co-activator of the glucocorticoid receptor | 9388192 | 17 NC_000017.10 (40465342..40540586, complement) |
| **STAT5 (STAT5A, STAT5B)** | GR can act as transcriptional coactivator for Stat5 and enhance Stat5-dependent transcription | 8878484 | 17 NC_000017.10 (40439565..40463961)<br><br>17 NC_000017.10 (40351195..40428478, complement) |
| **STAT6** | Physically and functionally interacts with GR in T-lymphocytes | 11150515 | 12 NC_000012.11 (57489187..57505196, complement) |
| **Thioredoxin(Trx)** | Thioredoxin negatively modulates GR function | 8958209 | 9 NC_000009.11 (113006092..113018920, complement) |
| **Mitochondrial Thioredoxin (Trx2)** | Mitochondrial thioredoxin has a regulatory role in GR and NFκB signaling pathways. Specifically, Trx2 stimulates the TNFα- | 19570036 | 22 NC_000022.10 (36863083..36878072, complement) |

| | | | |
|---|---|---|---|
| | induced NFκB activation and DEX-induced GR activation of reporter genes | | |
| **Thioredoxin reductase 1 (TrxR1)** | Overexpression of TrxR1 increases GR activity in specific cells | 17382897 | 12 NC_000012.11 (104609537..104744085) |
| **TRIM28** | TRIM28 enhances GR-regulated expression | 9742105 | 19 NC_000019.9 (59055824..59062087) |
| **NCOR1** | Represses the GR gene trough a GR-NCOR1-HDAC3 repression complex | 23428870 | 17 NC_000017.10 (15933408..16118874, complement) |
| **HDAC3** | Represses the GR gene through a GR-NCOR1-HDAC3 repression complex | 23428870 | 5 NC_000005.9 (141000443..141016423, complement) |
| **HDAC6** | Regulates GR signaling through HSP90 acetylation control | 22457490 | X NC_000023.10 (48660086..48683408) |
| **NR2F2** | NR2F2 represses the GR-stimulated transcriptional activity by tethering corepressors such as NCOR2 and NCOR1. GR stimulates NR2F2 transactivating factors. | 15265774 | 15 NC_000015.9 (96869157..96883492) |
| **NCOR2** | Partakes in NR2F2-dependent GR suppression | 15265774 | 12 NC_000012.11 (124808957..125052079, complement) |
| **NFKBIA** | GR activates its' specific gene to repress NFκB expression | 11694573 | 14 NC_000014.8 (35870716..35873960, complement) |
| **EGFR** | GR modulates EGFR function | 31052457 | 7 NC_000007.13 (55086678..55279262) |
| **HMGB1** | GR modulates HMGB1 expression | 21737101 | 13 NC_000013.10 (31032877..31191942, complement) |
| **RPS6KA5(MSK1)** | Liganded GR interacts with activated RPS6KA5 resulting in redistribution of a part of the nuclear RPSK6KA5 pool to the cytoplasm | 20456998 | 14 NC_000014.8 (91335086..91526993, complement) |
| **Casein kinase 2 (CSNK2A1, CSNK2A2, CSNK2B)** | It phosphorylates the Glucocorticoid Receptor | 23953592 | 20 NC_000020.10 (463338..524482, complement)<br><br>16 NC_000016.9 (58191811..58231782, complement)<br><br>6 NC_000006.11 (31632995..31637844) |

| | | | |
|---|---|---|---|
| **NLRP3** | GR binds on its' specific gene and regulates its' expression | 21940629 | 1 NC_000001.10 (247579247..247612410) |
| **Mcl-1** | GR directly binds on its' gene and regulates Mcl-1 expression | 20156337 | 1 NC_000001.10 (150547027..150552214, complement) |
| **NOXA** | GR directly binds on its' gene and regulates NOXA expression | 20156337 | 18 NC_000018.9 (57567153..57571538) |
| **KLF13** | GR binds on the KLF13 promoter to trigger its' expression | 25336632 | 15 NC_000015.9 (31619083..31670102) |
| **BIM** | Shows an intronic binding site for GR, that is activated in case of dexamethasone sensitivity | 25336632 | 2 NC_000002.11 (111878491..111926022) |
| **FOXO3** | GR induces its' transcription | 22848740 | 6 NC_000006.11 (108881026..109005972) |
| **BAK** | BAK co-precipitates with GR upon dexamethasone treatment | 27888447 | 6 NC_000006.11 (33540323..33548072, complement) |
| **Bcl-xL** | Bcl-xL co-precipitates with GR upon dexamethasone treatment | 27888447 | 20 NC_000020.10 (30252261..30311752, complement) |
| **PI3K (p85 subunit /PIK3R1-6)** | Physically interacts with GR, they, then, regulate the tlr2 signaling cascade | 19874421 | 5 NC_000005.9 (67511584..67597649)<br><br>19 NC_000019.9 (18263988..18281343<br><br>1 NC_000001.10 (46505812..46642167, complement)<br><br>3 NC_000003.11 (130397778..130465696, complement)<br><br>17 NC_000017.10 (8782233..8869029, complement)<br><br>17 NC_000017.10 (8706055..8770994, complement) |
| **Annexin1** | GR induces its' specific gene (ANXA1) | 16236742 | 9 NC_000009.11 (75766721..75785309) |
| **TSLP** | GR negatively regulates TSLP's gene expression | 23222642 | 5 NC_000005.9 (110405778..110413722) |

| | | | |
|---|---|---|---|
| **ST13** | ST13 promotes the functional maturation of GR | 27169854 | 22 NC_000022.10 (41220539..41253012, complement) |
| **PPID** | PPID anchors on the GR complex | 27169854 | 4 NC_000004.11 (159630279..159644552, complement) |
| **IRF8** | Its' gene is regulated by GR | 17185395 | 16 NC_000016.9 (85932774..85956212) |
| **LAD1** | Its' gene is regulated by GR | 17185395 | 1 NC_000001.10 (201349966..201368669, complement) |
| **IGFBP-1** | Its' gene is regulated by GR | 17185395 | 7 NC_000007.13 (45927959..45933267) |
| **PKAc (PRKACA PRKACB PRKACG)** | GR cross-couples with the catalytic subunit of PKA | 27169854 | 19 NC_000019.9 (14202500..14228559, complement)<br><br>1 NC_000001.10 (84543745..84704181)<br><br>9 NC_000009.11 (71627426..71635600, complement) |
| **TRIP6** | TRIP6 creates a complex with GR, which partakes in the receptor's transrepression ability | 27169854 | 7 NC_000007.13 (100464950..100471076) |
| **14-3-3 (14-3-3σ 14-3-3η 14-3-3ζ/δ)** | Takes part in a complex which features GR and Raf-1 | 27169854 | 1 NC_000001.10 (27189633..27190947)<br><br>22 NC_000022.10 (32340479..32353590)<br><br>8 NC_000008.10 (101930804..101965717, complement) |
| **Raf-1** | Takes part in a complex which features GR and 14-3-3 | 27169854 | 3 NC_000003.11 (12625100..12705700, complement) |
| **PPARγ** | Interacts with GR | 27169854 | 3 NC_000003.11 (12329349..12475855) |
| **PPARα** | Interacts with GR and they both act as immunosuppressors | 27169854 | 22 NC_000022.10 (46546458..46639653) |
| **LXR(α,β)** | LXR has both synergistic and opposing effects on GR | 27169854 | 11 NC_000011.9 (47269851..47290584)<br><br>19 NC_000019.9 (50879680..50886285) |

| | | | |
|---|---|---|---|
| **RARα & RXRα** | They both bind on GR and enhance its' transcriptional activity | 27169854 | 17 NC_000017.10 (38465423..38513895)<br><br>9 NC_000009.11 (137218316..137332431) |
| **Progesterone Receptor** | They possibly interact to repress IL-1β-driven COX-2 activation | 27169854 | 11 NC_000011.9 (100900355..101000544, complement) |
| **Androgen Receptor** | It creates a heterodimer with GR at a common DNA site, leading to mutual transcriptional inhibition | 27169854 | X NC_000023.10 (66763874..66950461) |
| **Estrogen Receptor alpha** | Its interaction with GR can have cooperative or antagonistic action on E2-regulated genes | 27169854 | 6 NC_000006.11 (152011631..152424409) |
| **Nur77** | Through protein-protein interaction GR antagonizes Nur77-depenent transcription on the Nur77 response element of the pomc gene | 27169854 | 12 NC_000012.11 (52416616..52453291) |
| **Dax-1** | Dax-1 competes coactivator NCOA2 in binding to the GR | 27169854 | X NC_000023.10 (30322539..30327495, complement) |
| **SOCS1** | GR ad SOCS1 create an intracellular complex and GCs increase the nuclear levels of SOCS1 | 18524780 | 16 NC_000016.9 (11348274..11350039, complement) |
| **Tbx21/T-bet** | GR interacts with Tbx21 and inhibits Tbx21's action | 27169854 | 17 NC_000017.10 (45810610..45823485) |
| **OGT** | OGT interacts with ligand-bound GR and potentiates the GR transreppression pathway | 27169854 | X NC_000023.10 (70752912..70795747) |
| **FOXA3** | FOXA3 mediates GR function in adipose tissue | 26957608 | 19 NC_000019.9 (46367518..46377055) |
| **PER2** | GR regulates its' function | 19805059 | 2 NC_000002.11 (239152679..239198678, complement) |
| **RSUME** | Possibly interacts with GR and takes part in the receptor's sumoylation | 27169854 | 1 NC_000001.10 (95699711..95712781) |
| **SUMO4** | SUMO4-induced GR sumoylation enhances GR DNA binding activity | 27169854 | 6 NC_000006.11 (149721284..149722182) |
| **Ubch7** | It interacts with GR and its' effects on the receptor depend on the cell culture studied | 27169854 | 22 NC_000022.10 (21903736..21978323) |
| **E6-AP** | E6-AP regulates GR transactivation | 27169854 | 15 NC_000015.9 (25582394..25684190, complement) |

**Supplementary Table 2.** SNPs of interest and polymorphisms, diseases, chemicals, and variants that commonly co-occur with mentioned SNPs in the same sentence.

| SNPs | Pub. | Variants Co. | Located in Gene | Diseases Co. | Chemicals Co. |
|---|---|---|---|---|---|
| rs4713916 | 49 | rs1360780<br>rs3800373<br>rs9470080<br>rs9296158<br>rs41423247<br>rs4713902<br>rs7997012<br>rs9394309<br>rs6265 | FKBP5 | -Depressive Disorder<br>-Major Depressive Disorder<br>-Wounds and Injuries<br>-Abusive Dwarfism Syndrome<br>-Anxiety Disorders | -hydrocortisone<br>-Citalopram<br>-Serotonin<br>-Ethanol<br>-C081489<br>-Steroids |
| rs1360780 | 271 | rs3800373<br>rs9296158<br>rs9470080<br>rs4713916<br>rs6265 | FKBP5 | -Depressive Disorder<br>-Post Traumatic Stress Disorder<br>-Wounds and Injuries<br>-Major Depressive Disorder<br>-Mental Disorders | -Hydrocortisone<br>-Ethanol<br>-Dexamethazone<br>-Tacrolimus |
| rs1045642 | 1984 | rs1128503<br>rs2032582<br>c.2677G>T,A<br>rs776746<br>rs2231142<br>rs4244285<br>rs1801133<br>rs717620<br>rs4149056 | ABCB1 | -Epilepsy<br>-Breast Neoplasms,<br>-Neoplasms<br>-Drug-Related Side Effects and Adverse Reactions<br>-Colorectal Neoplasms | -Clopidogrel<br>-Tacrolimus<br>-Cyclosporine<br>-Digoxin<br>-Methotrexate<br>-Peptide T amide |
| rs3842 | 26 | rs1045642<br>rs3745274<br>rs776746<br>rs1128503<br>rs10264272 | ABCB1 | -Hypertension<br>-Diabetes Mellitus<br>-Lung Neoplasms<br>-Dyslipidemias<br>-HIV Infections | -Efavirenz<br>-1-chloro-2-hydroxy-3-butene<br>-Carbon<br>-1,7,9,11-tetrahydroxy-3-methyl-8,13-dioxo-5,6,8,13-tetrahydrobenzo(a)tetracene-2-carboxylic acid |
| rs1922242 | 8 | rs1045642<br>rs1202184<br>rs10808072<br>rs3213619<br>rs1128503<br>rs2032582<br>rs1202168 | ABCB1 | -Renal Cell Carcinoma<br>-Depressive Disorder<br>-Anxiety Disorders<br>-Seizures | -12-(4'-azido-2'-nitrophenoxy)dodecanoyl-coenzyme A<br>-Oxygen<br>-Thulium |
| rs2235046 | 9 | rs1128503<br>rs10276036<br>rs1202169<br>rs4148738<br>rs1045642 | ABCB1 | -Renal Insufficiency<br>-Zellweger Syndrome<br>-Lung Neoplasms<br>-N syndrome<br>-Bilateral Multicystic Renal Dysplasia | -C554682<br>-Apixaban<br>-C065179<br>-Nitrogen<br>-C503223<br>-Interleukin-2 Receptor beta Subunit<br>-Carbon |
| rs2235013 | 8 | rs1045642<br>rs1128503<br>rs2235033 | ABCB1 | -Follicular Thyroid Cancer<br>-Lung Neoplasms<br>-Proteinuria | -Cyclosporine<br>-Pentalysine |

| | | | | | |
|---|---|---|---|---|---|
| | | rs2032582<br>rs1202179<br>rs1695<br>rs10276036<br>rs2235046<br>rs9282564 | | -Ataxia Telangiectasia<br>-Zellweger Syndrome | -Alanyl-alanyl-alanyl-alanine<br>-Methionylmethionine<br>-Seryl-seryl-seryl-arginine<br>-Leucylleucine<br>-Peptide T amide<br>-2'-deoxy-5-fluoro-3'-thiacytidine<br>-Carbon<br>-Interleukin-2 Receptor beta Subunit |
| rs2235035 | 6 | rs1202169<br>rs2032582<br>rs1045642<br>rs1138272<br>rs4520<br>rs1292798<br>rs4891<br>rs1027649<br>rs2235046 | ABCB1 | -N syndrome<br>-Ataxia Telangiectasia | -Alanyl-alanyl-alanyl-alanine<br>-Triglycerides<br>-Angoletin<br>-Seryl-seryl-seryl-arginine<br>-Peptide T amide |
| rs2235033 | 6 | rs1128503<br>rs1045642<br>rs2235013<br>rs2235046<br>rs4148738<br>rs4680<br>rs2273697<br>rs10276036<br>rs4437575 | ABCB1 | -Zellweger Syndrome | -Sulfur<br>-Carbon<br>-Interleukin-2 Receptor beta Subunit<br>-Daunorubicin<br>-Cyclosporine<br>-Daunorubicinol |
| rs10276036 | 15 | rs1128503<br>rs2235046<br>rs1202169<br>rs1202167<br>rs1202168<br>rs4148738 | ABCB1 | -Neutropenia<br>-Diabetes Mellitus<br>-Hypertension<br>-Breast Neoplasms<br>-Neoplasms | -C554682<br>-Doxorubicin<br>-Glycyl-glycyl-sarcosine<br>-Warfarin<br>-Serotonin<br>-Apixaban<br>-Irinotecan<br>-Superoxides<br>-Adenosine triphosphate<br>-Guanosine |
| rs1922240 | 2 | rs6591256<br>rs1338062<br>rs754814<br>rs7793196<br>rs7223183<br>rs11869640<br>rs4148732<br>rs7299040<br>rs5993875 | ABCB1 | -Pain<br>-N-Syndrome | -Morphine |
| rs868755 | 9 | rs1858923<br>rs1202168<br>rs1045642<br>rs10280623<br>rs4148738<br>rs2032582<br>rs7779562 | ABCB1 | -Drug-Related Side Effects and Adverse Reactions<br>- Zellweger Syndrome<br>- Colorectal Neoplasms | -Carbon<br>-Interleukin-2 Receptor beta Subunit |

| | | | | | |
|---|---|---|---|---|---|
| | | rs10808072<br>rs2235048 | | | |
| rs13237132 | 2 | rs2235023<br>rs4148732<br>rs12334183<br>rs10264990<br>rs238416<br>rs12129768<br>rs11188148<br>rs1678607<br>rs50872 | ABCB1 | - Ovarian Neoplasms<br>- Bradycardia<br>-Von Hippel-Lindau Disease | - |
| rs1202170 | 1 | rs1799971<br>rs2235033<br>rs3024971<br>rs3786047<br>rs2235013<br>rs167769<br>rs1045642<br>rs1045280<br>rs2036657 | ABCB1 | - | - |
| rs1202168 | 11 | rs1045642<br>rs1128503<br>rs2032582<br>rs868755<br>rs1202169<br>rs2235046<br>rs10276036<br>rs4148738<br>rs1202167 | ABCB1 | - Colorectal Neoplasms<br>- Zellweger Syndrome<br>-Neonatal Hyperbilirubinemia | -C554682<br>-Thulium<br>-C065179<br>-Apixaban<br>-12-(4'-azido-2'-nitrophenoxy)dodecanoyl-coenzyme A<br>-Interleukin-2 Receptor beta Subunit<br>-Carbon<br>-Oxygen<br>-C503223 |
| rs1016793 | 2 | rs6961665<br>rs41277128<br>rs62578960<br>rs10985911<br>rs1202168<br>rs116855710<br>rs114717568<br>rs11673270<br>rs11083571 | ABCB1 | -Zellweger Syndrome | -Carbon<br>-Interleukin-2 Receptor beta Subunit |
| rs2235018 | 1 | rs34800935<br>rs7793933<br>rs2188526<br>rs1045642<br>rs6949448<br>rs1922244<br>rs2235048<br>rs4148738<br>rs7787082<br>rs12720464 | ABCB1 | - | -Carbon |
| rs1211152 | 3 | rs1045642<br>rs10264990<br>rs1202184<br>rs17327624<br>rs6946119 | ABCB1 | - | - |
| rs2235074 | 2 | rs35979566<br>rs2279342<br>rs1202169 | ABCB1 | -Myelodysplastic Syndromes | -Adenosine Triphosphate<br>-Nitrogen |

| | | | | | |
|---|---|---|---|---|---|
| | | rs6591722<br>rs1010570<br>rs4244285<br>rs4148329<br>rs1042838<br>rs7801671 | | | |
| rs2214102 | 12 | rs1045642<br>rs1128503<br>rs2229109<br>rs2032582<br>rs3213619<br>rs9282564 | ABCB1 | -Breast Neoplasms<br>-Ataxia Telangiectasia | -Decaglycine<br>-Peptide T amide<br>-His-His-His-His-His-His<br>-Seryl-seryl-seryl-arginine<br>-Leucylleucine<br>-Triamcinolone<br>-Progestins<br>-Diprotin A<br>-Asparagyl-aspargyl-tryptophyl-asparagyl-asparagine<br>-1,7,9,11-tetrahydroxy-3-methyl-8,13-dioxo-5,6,8,13-tetrahydrobenzo(a)tetracene-2-carboxylic acid |
| rs3213619 | 75 | rs1045642<br>rs1128503<br>rs2032582<br>c.2677G>T,A<br>rs776746 | ABCB1 | -Colorectal Neoplasms<br>-Hypertension<br>-Drug-Related Side Effects and Adverse Reactions<br>-Dyslipidemias<br>-Diabetes Mellitus | -Tacrolimus<br>-Paclitaxel<br>-Docetaxel<br>-C097613<br>-Vasoactive intestinal constrictor<br>-Taxane<br>-Methotrexate-alpha-phenylalanine<br>-Carbon<br>-Decaglycine<br>-Cyclosporine |
| rs4986893 | 276 | rs4244285<br>rs12248560<br>rs1057910<br>rs1799853<br>rs1045642 | CYP2C19 | -Drug-Related Side Effects and Adverse Reactions<br>-Hypertension<br>-Breast Neoplasms<br>-Thrombosis<br>-Stroke | -Clopidogrel<br>-Warfarin<br>-Simvastatin<br>-Carbon<br>-Tryptophyl-arginyl-tryptophyl-tryptophyl-tryptophyl-tryptophanamide |
| rs4244285 | 475 | rs4986893<br>rs12248560<br>rs1057910<br>rs1799853<br>rs1045642 | CYP2C19 | -Drug-Related Side Effects and Adverse Reactions<br>-Blood Platelet Disorders<br>-Breast Neoplasms<br>-Hemorrhage<br>-Thrombosis | Clopidogrel<br>Warfarin<br>Simvastatin<br>Aspirin<br>Carbon |
| rs72552267 | 31 | rs41291556<br>rs28399504<br>rs4986893<br>rs56337013 | CYP2C19 | -N syndrome<br>-Abnormal Reflex<br>-Norrie Disease | -Clopidogrel<br>-Warfarin<br>-Simvastatin<br>-Metformin |

| | | | | | |
|---|---|---|---|---|---|
| | | rs4244285 | | -Drug-Related Side Effects and Adverse Reactions<br>-Acute Coronary Syndrome | -Tacrolimus |
| rs1057910 | 512 | rs1799853<br>rs9923231<br>rs4244285<br>rs2108622<br>rs4986893 | CYP2C9 | -Drug-Related Side Effects and Adverse Reactions<br>-Hemorrhage<br>-Diabetes Mellitus<br>-Neoplasms<br>-Hypertension | -Warfarin<br>-Clopidogrel<br>-Simvastatin<br>-Ciproxifan<br>-Phenytoin |
| rs7089580 | 12 | rs61162043<br>rs9923231<br>rs1799853<br>rs7900194<br>rs28371686<br>rs28371685<br>rs1057910 | CYP2C9 | -Atrial Fibrillation<br>-Norrie Disease | -Warfarin<br>-Amiodarone<br>-Carcinoma-associated --Antigen 17-1A<br>-Synthetic SNP-1 protein |
| rs4917639 | 18 | rs1057910<br>rs9923231<br>rs1799853<br>rs7294<br>rs10871454 | CYP2C9 | -Stroke<br>-Zellweger Syndrome<br>-Intracranial Hemorrhages | -Warfarin<br>-C065179<br>-Carvedilol<br>-S-imvastatin<br>-Acenocoumarol<br>-Metoprolol<br>-Pravastatin<br>-Vitamin K1 oxide<br>-Clopidogrel<br>-Sodium |
| rs4646450 | 12 | rs15524<br>rs776746<br>rs4244285<br>rs1128503<br>rs1045642 | CYP3A5 | -Cardiotoxicity<br>-Urinary Bladder Neoplasms<br>-Graft vs Host Disease | -12-(4'-azido-2'-nitrophenoxy)dodecanoyl-coenzyme A<br>-Tacrolimus<br>-Aldrin<br>-Lopinavir<br>-Sch 601324<br>-Alachlor<br>-Cyanazine<br>-Calcium<br>-Poly(acrylamide-co-crotonic acid)<br>-Dehydroepiandrosterone Sulfate |
| rs776746 | 524 | rs1045642<br>rs2740574<br>rs1128503<br>rs2032582<br>rs35599367 | CYP3A5 | -Drug-Related Side Effects and Adverse Reactions<br>-Hypertension<br>-Neoplasms<br>-Diabetes Mellitus<br>-Non-Small-Cell Lung Carcinoma | -Tacrolimus<br>-Simvastatin<br>-Cyclosporine<br>-Clopidogrel<br>-Warfarin<br>-Sunitinib |
| rs1043618 | 59 | rs2227956<br>rs1061581<br>rs1008438<br>rs2075800<br>rs2763979 | HSPA1A | -Major Depressive Disorder<br>-Alzheimer Disease<br>-Depressive Disorder<br>-Glaucoma<br>-Neoplasms | -2-carboxyarabinitol 1-phosphate<br>Gastrofenzin<br>Poly-aluminum-chloride-sulfate<br>Nitroglycerin<br>Glycyl-threonine<br>Oxytocin, Glu(4)-<br>Hydrogen<br>Nitrogen<br>Tyrosyl-lysine |

| | | | | | |
|---|---|---|---|---|---|
| | | | | | -Human LRRN2 protein |
| rs6457452 | 11 | rs2763979<br>rs1061581<br>rs17200983<br>rs13118<br>rs150142878<br>rs9267546<br>rs11538264<br>rs9267547<br>rs4576240 | HSPA1B | -Alopecia Areata<br>-Schizophrenia<br>-Paranoid Schizophrenia<br>-Anemia<br>-Malaria | -15-hydroxy-5,8,11,13-eicosatetraenoic acid<br>-3,4,5-trichloroguaiacol<br>Cholesterol<br>-Methacholine Chloride<br>-Triglycerides<br>-Prostaglandins<br>-Uric Acid<br>-Aspirin<br>-Carbon |
| rs2227956 | 63 | rs1061581<br>rs1043618<br>rs2075800<br>rs2763979<br>rs662 | HSPA1L | -Stomach Neoplasms<br>-Male Infertility<br>-Neoplasms<br>-Ataxia Telangiectasia<br>-Diabetic Foot | -Peptide T amide<br>-Methionylmethionine<br>-Glycyl-threonine<br>-Methionine<br>-1,10-phenanthroline-5,6-dione |
| rs2227955 | 2 | rs2227956<br>rs2075800<br>rs35326839<br>rs10117<br>rs116768554<br>rs14355<br>rs566393477<br>rs1042881<br>rs34620296 | HSPA1L | -Ataxia Telangiectasia | -Alanine<br>-Glycine<br>-1,10-phenanthroline-5,6--dione<br>-Peptide T amide<br>-Glycyl-threonine<br>-Methionine<br>-Arginyl-glutamine |
| rs34620296 | 2 | rs139193421<br>p.K73S<br>rs2075799<br>rs368138379<br>rs199780750<br>rs139868987<br>rs2227956<br>c.515_517del<br>c.218A>G | HSPA1L | -Multiple Hamartoma Syndrome<br>-Crohn Disease<br>-Proctitis<br>-Colitis<br>-Gastritis | -1,10-phenanthroline-5,6-dione<br>-IS 23<br>-Peptide T amide |
| rs368138379 | 1 | rs199780750<br>rs2075799<br>rs116768554<br>rs1460318497<br>rs566393477<br>rs35326839<br>rs2075800<br>p.172del<br>rs35347921<br>rs9469057 | HSPA1L | -Multiple Hamartoma Syndrome<br>-Crohn Disease<br>-Proctitis<br>-Colitis<br>-Gastritis | -1,10-phenanthroline-5,6-dione<br>-Peptide T amide |
| rs2279744 | 373 | rs1042522<br>rs117039649<br>rs1801270<br>rs25487<br>rs9344 | MDM2 | -Neoplasms<br>-Lung Neoplasms<br>-Breast Neoplasms<br>-Stomach Neoplasms<br>-Endometrial Neoplasms | Arginylarginine<br>Estrogens<br>Synthetic SNP-1 protein<br>Gastrofenzin<br>Nitrogen<br>Cisplatin |

| | | | | | |
|---|---|---|---|---|---|
| rs2228570 | 717 | rs1544410<br>rs731236<br>rs7975232<br>rs11568820<br>rs7041 | VDR | -Ovarian Neoplasms -Asthma<br>-Breast Neoplasms<br>-Neoplasms<br>-Multiple Sclerosis | -Vitamin D<br>-25-hydroxyvitamin D3-bromoacetate<br>-Calcium<br>-Poly If<br>-Peptide T amide |
| rs201753350 | 18 | rs28934576<br>rs1042522<br>rs1800371<br>rs1800370<br>rs730882025<br>rs28934578<br>rs1800372<br>rs104886003<br>rs1057519991 | TP53 | -Rhabdomyosarcoma<br>-Emanuel Syndrome<br>-Acute Myeloid Leukemia<br>-Neoplasms<br>-Norrie Disease | -AT 61<br>-Arginyl-tryptophyl-arginine<br>Nitrogen<br>Histocompatibility Antigen H-2D<br>Leucylleucine<br>-3-bromoacetoxyandrostan-17-one<br>-2-(3,4-dimethoxyphenyl)-5-amino-2-isopropylvaleronitrile<br>-chromozym TH<br>-(arginine)9-cysteinyl-glutaminyl-cysteinyl-arginyl-arginyl-lysyl-asparagine<br>-H 189 |
| rs1800206 | 217 | rs2016520<br>rs1801282<br>rs4253778<br>rs3856806<br>rs135539<br>rs1805192 | PPARA | -Diabetes Mellitus<br>-Obesity<br>-Metabolic Diseases<br>-Type 2 Diabetes Mellitus<br>-Atherosclerosis | -Triglycerides<br>- Omega-3 Fatty Acids<br>-Fatty Acids<br>-Cholesterol<br>-Unsaturated Fatty Acids |
| rs1801282 | 979 | rs7903146<br>rs5219<br>rs13266634<br>rs4402960<br>rs10811661<br>rs1111875<br>rs3856806<br>rs864745<br>rs7961581 | PPARG | -Diabetes Mellitus<br>-Obesity<br>-Type 2 Diabetes Mellitus<br>-Metabolic Diseases<br>-Neoplasms<br>-Hypertension<br>-Insulin Resistance<br>-Coronary Disease<br>-Colorectal Neoplasms<br>-Polycystic Ovary Syndrome | -Alanyl-alanyl-alanyl-alanine<br>-N-nitroso-prolylalanine<br>-Glucose<br>-Troglitazone<br>-Thiazolidinediones<br>-Cholesterol<br>-Triglycerides<br>-Ethanol<br>-Potassium<br>-Fatty Acids |
| rs28936407 | 6 | rs121909242<br>rs72551362<br>rs121909243<br>rs72551364<br>rs72551363<br>rs121909244 | PPARG | -Neoplasms<br>-Colorectal Neoplasms<br>-Lipodystrophy<br>-Lipid Metabolism Disorders<br>-Migraine Disorders | -Urea<br>-Hydrogen<br>-Rosiglitazone<br>-Lecithin emulsion safflower oil |
| rs237025 | 132 | rs2476601<br>rs577001<br>rs1805010<br>rs237024 | SUMO4 | -Diabetes Mellitus<br>-Type 1 Diabetes Mellitus<br>-Type 3 Axenfeld-Rieger syndrome | -Methionine<br>-Valine-valine-saquinavir<br>Tacrolimus |

| | | | | | |
|---|---|---|---|---|---|
| | | rs1800872<br>rs2243250 | | -Type 2 Diabetes Mellitus<br>-Diabetic Nephropathies | -(Z)-2-amino-5-chlorobenzophenonamidinohydrazone acetate<br>-Nitrogen<br>-Triglycerides |
| rs118203914 | 1 | p.E411X (1)<br>rs761817519 (1)<br>p.L201R (1)<br>rs758306831 (1)<br>rs775488556 (1) | TAT | Type 2C Congenital Disorder of Glycosylation | |
